# Nup93 and CTCF co-modulate spatiotemporal dynamics and function of the HOXA gene cluster during differentiation

**DOI:** 10.1101/646224

**Authors:** Ajay S. Labade, Adwait Salvi, Krishanpal Karmodiya, Kundan Sengupta

## Abstract

Nucleoporins regulate nuclear transport. In addition, nucleoporins also modulate chromatin organization and gene expression. Here we investigated the role of nucleoporin Nup93, in regulating HOXA gene expression during differentiation. ChIP-Seq analysis revealed that Nup93 associates with genes involved in development and differentiation. Furthermore, Nup93 occupancy significantly overlaps with CTCF. Interestingly, Nup93 and CTCF show antagonistic roles in regulating 3’ and 5’ end HOXA genes in undifferentiated cells. The HOXA gene locus untethered from the nuclear periphery upon Nup93 but not CTCF depletion, consistent with its upregulation. Remarkably, occupancy of Nup93 and CTCF on HOXA gene locus progressively declined during differentiation but was restored in differentiated cells, consistent with the rerepression and re-localization of the HOXA gene locus with the nuclear periphery upon differentiation. In summary, Nup93 is a key modulator of the spatiotemporal dynamics and function of the HOXA gene locus during differentiation.

## INTRODUCTION

The nuclear pore complex (NPC) consists of 30 different subunits forming a nuclear-cytoplasmic transport channel (D’Angelo and Hetzer 2008). In addition to nuclear transport, nucleoporins regulate gene expression by associating with chromatin (Capelson and Hetzer 2009; Capelson et al. 2010; Kalverda et al. 2010; Vaquerizas et al. 2010; Sood and Brickner 2014; Ibarra et al. 2016). Based on their stability and location at the nuclear membrane, nucleoporins are classified as (i) scaffold nucleoporins - a highly stable core ring-like structure of the NPC and (ii) peripheral nucleoporins, which include the central channel, nuclear basket and cytoplasmic filaments (D’Angelo and Hetzer 2008).

In mammals, mechanisms of nucleoporin mediated gene expression are unclear. However, emerging evidence from model organisms such as *Drosophila, C.elegans*, and yeast, suggest that nucleoporins also regulate gene expression (Griffis et al. 2002; Schmid et al. 2006; Capelson et al. 2010; Ikegami and Lieb 2013; Rohner et al. 2013; Pascual-Garcia et al. 2017). ChIP-chip (Chromatin immunoprecipitation followed by tiling microarray) revealed that nucleoporin Nup93 associates with human chromosomes 5, 7 and 16 in HeLa cells (Brown et al. 2008). Dam ID analyses of genome-wide interaction of Nup93 revealed its association with super-enhancers, that regulate expression of cell identity genes (Ibarra et al. 2016). Nucleoporin - Nup210, shows tissue-specific expression and is involved in muscle cell differentiation, maturation and survival of differentiated cells (D’Angelo et al. 2012; Raices et al. 2017). Nup50 depletion inhibits differentiation of C2C12 cells into myotubes (Buchwalter et al. 2014). The mobile nucleoporin Nup98 (a phenylalanine-glycine (FG) repeat-containing nucleoporin), associates with genes involved in development and differentiation (Liang et al. 2013). The mobile Nup153, represses developmental genes by recruiting PRC1 complex at their promoters in mouse ES cells (Jacinto et al. 2015). Taken together, these findings highlight the role of nucleoporins in regulating expression of genes involved in development. However, nucleoporin function in regulating gene expression in a temporal manner during differentiation is unclear.

Nup93 is a relatively stable nucleoporin of the NPC (Daigle et al. 2001). Fluorescence Recovery After Photobleaching (FRAP) shows that Nup93 has a relatively long residence time at the NPC compared to other Nups with a low diffusion rate of K_off_: 4.0 ± 3.4 × 10^-6^ s^-1^ (Rabut et al. 2004). The high stability of Nup93, renders it as a chromatin anchor, potentially with heterochromatin, consistent with its relatively peripheral nuclear localization. ChIP-qPCR shows that Nup93 associates with the 3’ end of HOXA gene cluster and represses its expression in differentiated cells. The HOXA gene cluster (~150 Kb) maps to human chromosome 7, encodes eleven transcription factors and is expressed co-linearly during early stages of differentiation. HOXA genes (3’ end) are expressed early during differentiation, followed by expression of the 5’ end. Disruption of the temporal collinearity of HOXA gene expression during differentiation is associated with developmental defects (Aubin et al. 1997; Mallo and Alonso 2013). The untimely expression of HOXA genes is associated with cancers of the breast and lung (Novak et al. 2006; Bitu et al. 2012). Therefore, temporal and sequential silencing of HOXA gene expression is essential for normal development and differentiation (Mallo and Alonso 2013; Montavon and Soshnikova 2014). Chromatin conformation capture studies show that chromatin looping of the HOXA gene cluster mediated by CTCF, regulates HOXA gene expression during differentiation (Rousseau et al. 2014; Narendra et al. 2015). CTCF associates with multiple highly conserved CTCF binding sites (CBS) at the 5’-end of HOXA gene cluster, thereby regulating HOXA gene organization and expression (Rousseau et al. 2014; Xu et al. 2014). ChIP-Chip revealed that CTCF associated regions are enriched within Nup93 binding sites in HeLa cells (Brown et al. 2008). However, the crosstalk between the genome organizer CTCF, and Nup93 in regulating HOXA gene expression during differentiation remains elusive.

Here, we show that Nup93 and CTCF have antagonistic roles in regulating the spatiotemporal organization and function of HOXA gene locus during differentiation. ChIP-seq analyses of Nup93 revealed a considerable overlap between Nup93 and CTCF binding sites. Furthermore, Nup93 and CTCF depletion show antagonistic effects in modulating expression of HOXA genes in the 3’ and 5’ regions of the HOXA gene cluster. In addition, HOXA gene locus was repositioned away from the nuclear periphery upon activation but relocates proximal to the nuclear periphery, consistent with its attenuation. Remarkably, untethering of the HOXA locus from the nuclear periphery correlates with a reduced occupancy of Nup93 on HOXA. Furthermore, Nup93 and CTCF, re-occupy HOXA gene locus by Day 21 of differentiation. In summary, these results, reveal a dynamic interplay between Nup93 and CTCF in the regulation of HOXA gene expression during differentiation.

## RESULTS

### Nup93 occupancy is enriched on genes involved in development and differentiation

Chromatin immunoprecipitation of Nup93 followed by chromosome tiling array analyses in Hela cells revealed that Nup93 associates with the HOXA gene locus on human chromosome 7 (Brown et al. 2008). The Nup93 sub-complex associates with and represses HOXA gene cluster in differentiated cells (Labade et al. 2016). We examined the role of Nup93 in gene regulation by performing ChIP-Sequencing (ChIP-Seq) analyses of Nup93 in diploid colorectal cancer cells - DLD-1 (Supplemental Fig. S1A and B). ChIP-sequencing analyses revealed 404 high-confidence Nup93 binding sites in the human genome (Additional file 1). Furthermore, Nup93 was maximally enriched on intergenic (~46%), intronic (~34%) and promoter regions (~8%) (Fig. 1A). Nup93 was significantly enriched at Transcription Start Sites (TSS) of annotated genes (Supplemental Fig. S1C). Human Chromosome 2 showed the highest occupancy of Nup93 binding sites (~9.77%) (Supplemental Fig. S1D-F). While, gene-rich Chr.19 and gene poor chromosome 18, showed occupancy of ~5.51% and ~2% respectively (Supplemental Fig. S1D-F). Notwithstanding its relatively peripheral nuclear localization, human Chr.X, hardly showed enrichment of Nup93 binding sites (Supplemental Fig. S1D-F) (Chen et al. 2016). Furthermore, Nup93 binding did not reveal a correlation with either gene density or chromosome size (Supplemental Fig. S1E and F), implying that Nup93 occupancy may not correlate with the non-random organization of chromosome territories in the interphase nucleus (Hadlaczky et al. 1986; Zink et al. 1998). Taken together, Nup93 is enriched on promoter-proximal regions, consistent with its role in gene regulation (Brown et al. 2008; Ikegami and Lieb 2013; Ibarra et al. 2016).

**Figure 1.**
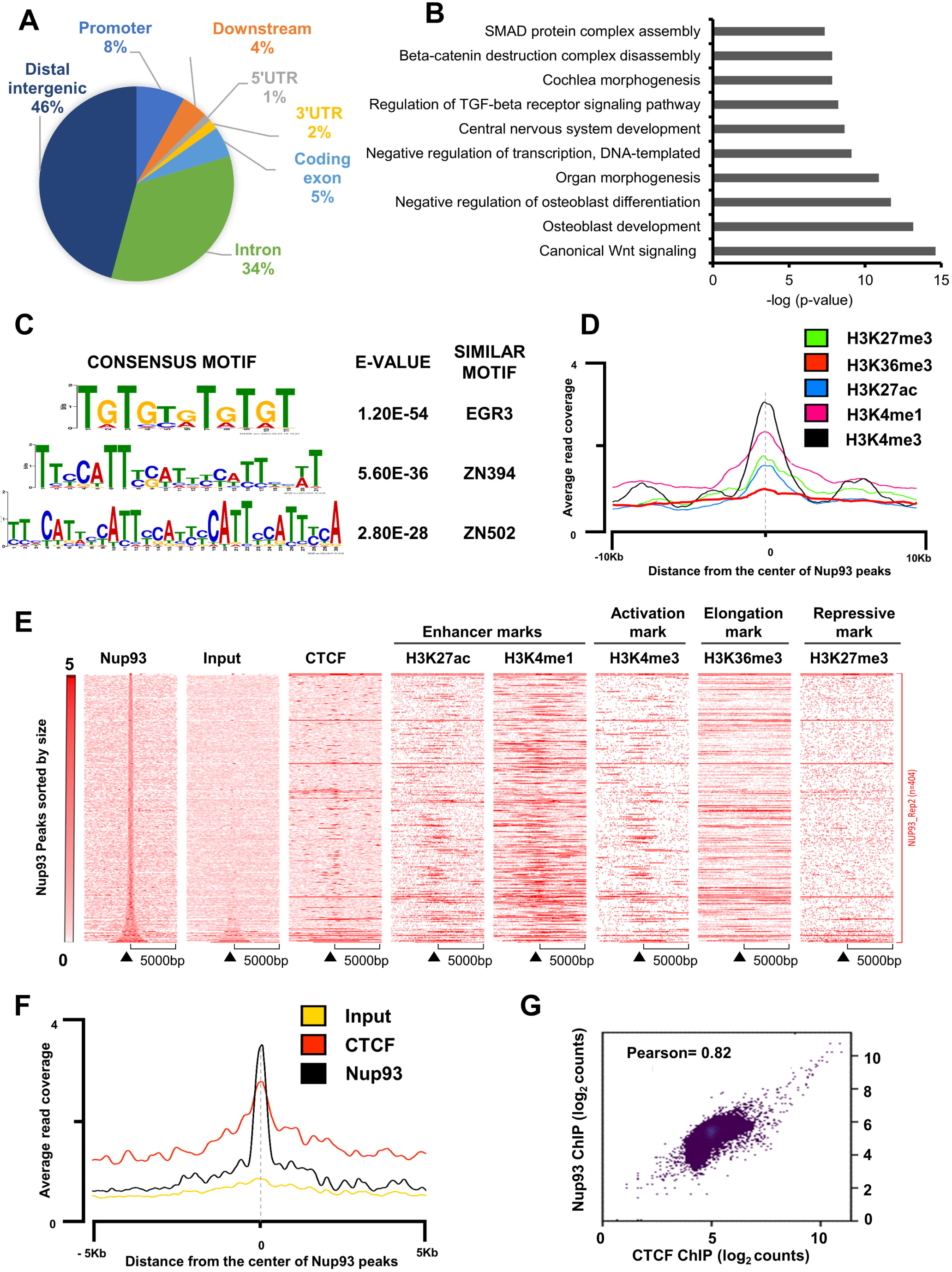
Genome-wide occupancy of Nup93. (A) Distribution of Nup93 binding peaks relative to various genomic elements [promoters (−3Kb from TSS), downstream region (+4Kb of TSS), 5’ UTR, 3’ UTR, introns and distal intergenic regions]. (B) Biological processes enriched for Nup93 associated genes as revealed by DAVID gene ontology analysis. (C) Top DNA-binding motifs enriched among binding sites of Nup93 as revealed by MEME-ChIP analysis, with corresponding p values. (D) Histogram showing an average distribution of tag density (intensity on Y-axis, tags/ million) of Repressive mark (H3K27me3), Elongation mark (H3K36me3), Enhancer marks (H3K27ac and H3K4me1) and Active mark (H3K4me3) mapped by ChIP-seq in DLD-1 cells relative to the center of Nup93 binding peaks. (E) Enrichment heatmaps of Nup93 ChIP-seq compared with Input, Nup93, CTCF, H3K27ac, H3K4me1, H3K4me3, H3K36me3, and H3K27me3 in DLD-1 cells. Peaks are sorted by size around the center of Nup93 binding peaks. (F) Histogram showing an average distribution of tag density (intensity on Y-axis, tags/million) of Input, Nup93 and CTCF reads mapped by ChIP-seq in DLD-1 cells relative to the center of Nup93 binding peaks. (G) Scatter plot showing Pearson’s correlation coefficient (r) between signal intensities (Log_2_ read counts in 10kb bins) of Nup93 and CTCF ChIP-seq data sets from DLD-1 cells. Nup93 occupancy positively correlates (r=0.82) with CTCF occupancy from DLD-1 cells.

We investigated into the functional role of Nup93, by performing Gene Ontology (GO) analyses using DAVID (Huang et al. 2009a, 2009b). Nup93 associated genes were enriched in canonical Wnt-signaling, osteoblast development, morphogenesis and central nervous system development (Fig. 1B). KEGG analyses (Kanehisa and Goto 2000) of Nup93 associated genes revealed pathways involved in development and differentiation such as (i) Hippo signaling (ii) Wnt-signaling and (iii) Signaling pathways regulating pluripotency of stem cells (Supplemental Fig. S2A). Nup93 bound sequences were enriched for consensus DNA motifs as that of transcription factors such as - EGR3, ZN394, ZN502, RREB1, MAZ and FOXG1 (Fig. 1C and Supplemental Fig. S2B), associated with transcriptional regulation of neuronal development, and neural differentiation (Eldredge et al. 2008; Quach et al. 2013; Wang et al. 2013; Bonomo et al. 2014). Taken together, these results implicate Nup93 in the regulation of genes associated with development and differentiation.

### Nup93 occupancy overlaps with CTCF binding sites

We determined the status of histone marks on Nup93 binding regions and analyzed the distribution of (i) active histone marks (H3K4me3) (ii) repressive mark (H3K27me3) (iii) enhancer marks (H3K27ac and H3K4me1) and (iv) elongation mark (H3K36me3) on ±5kb region from the center of Nup93 binding peaks (Fig. 1D and E). This showed a striking correlation between Nup93 binding sites with enhancer marks - H3K4me1 and H3K27ac (Fig. 1D and E), consistent with the enrichment of super-enhancer marks (H3K27ac) on Nup93 binding sites in U2OS cells (Ibarra et al. 2016). While the active histone mark H3K4me3 was enriched on Nup93 binding sites, the elongation mark (H3K36me3) was hardly enriched (Fig. 1D and E). Furthermore, transcription factor enrichment analysis using ReMap revealed that Nup93 binding sites overlap with that of CTCF (Supplementary Fig. S2C). Nup93 and CTCF occupancy compared from ChIP-Seq data sets of DLD-1 cells, revealed a significant enrichment of Nup93 peaks centered around CTCF binding sites (Fig. 1F). Nup93 and CTCF binding sites positively correlate with one another (Fig. 1G). CTCF preferentially binds to a conserved 11bp DNA motif [CCCTCGGTGGC] (Holohan et al. 2007). FIMO (Find Individual Motif Occurrences) analyses revealed 113 CTCF motif occurrences within Nup93 binding regions indicating that CTCF motifs were significantly enriched within Nup93 binding sites (Bailey et al. 2009). Of note, both Nup93 and CTCF were enriched on the HOXA1 promoter (Supplemental Fig. S2D). Taken together, Nup93 and CTCF are co-enriched at distal genomic regions such as super-enhancers and share common binding sites in the genome (Ibarra et al. 2016; Willi et al. 2017), further underscoring the role of Nup93 in regulating chromatin organization.

### Nup93 and CTCF levels are maintained during differentiation

ChIP-seq analyses revealed a potential crosstalk between Nup93 and CTCF in gene regulation. Nup93 and CTCF are enriched on super-enhancers associated with differentiation. In addition, Nup93 represses HOXA genes in differentiated cells (Labade et al. 2016), whereas CTCF regulates HOXA gene expression as it functions as an insulator demarcating the HOXA gene cluster. We determined the role of Nup93 and CTCF in HOXA gene regulation during differentiation. HOXA expression is activated upon retinoic acid (RA) treatment in NT2/D1 human embryonal teratocarcinoma cells that differentiate into neuronal cells - a well-established paradigm to study HOXA gene expression in a temporal manner (Simeone et al. 1990; Xu et al. 2014). We therefore, employed NT2/D1 cells as a model of cellular differentiation to investigate into the molecular mechanisms that regulate HOXA gene expression (Xu et al. 2014).

We treated NT2/D1 cells with RA (10μM) and followed their differentiation status for 8 Days, and up to 21 days respectively. NT2/D1 cells showed notable changes in their morphology accompanied by a progressive decrease in transcript levels of pluripotency genes - Oct4, Sox2, and Nanog (Fig. 2A-C). Furthermore, *Pax6* - a marker of neuronal differentiation, shows an increase followed by decrease in its levels during differentiation (Fig. 2A-C; Supplemental Fig. S3A-*Pax6* levels) (Bel-Vialar et al. 2007; Thakurela et al. 2016). In addition, FACS analyses revealed a marginal decrease in G2 sub-population, suggestive of reduced cell division during differentiation (Supplemental Fig. S3B and C). We next, performed qRT-PCR and immunoblotting across an 8 Day and 21 Day regimen of RA treatment, which did not show significant changes in either transcript or protein levels of Nup93 sub-complex (Nup93, Nup188, and Nup205) and CTCF (Fig. 2D and E; Supplemental Fig. S3D). This further underscores the stability of Nup93 and its interactors during differentiation.

**Figure 2.**
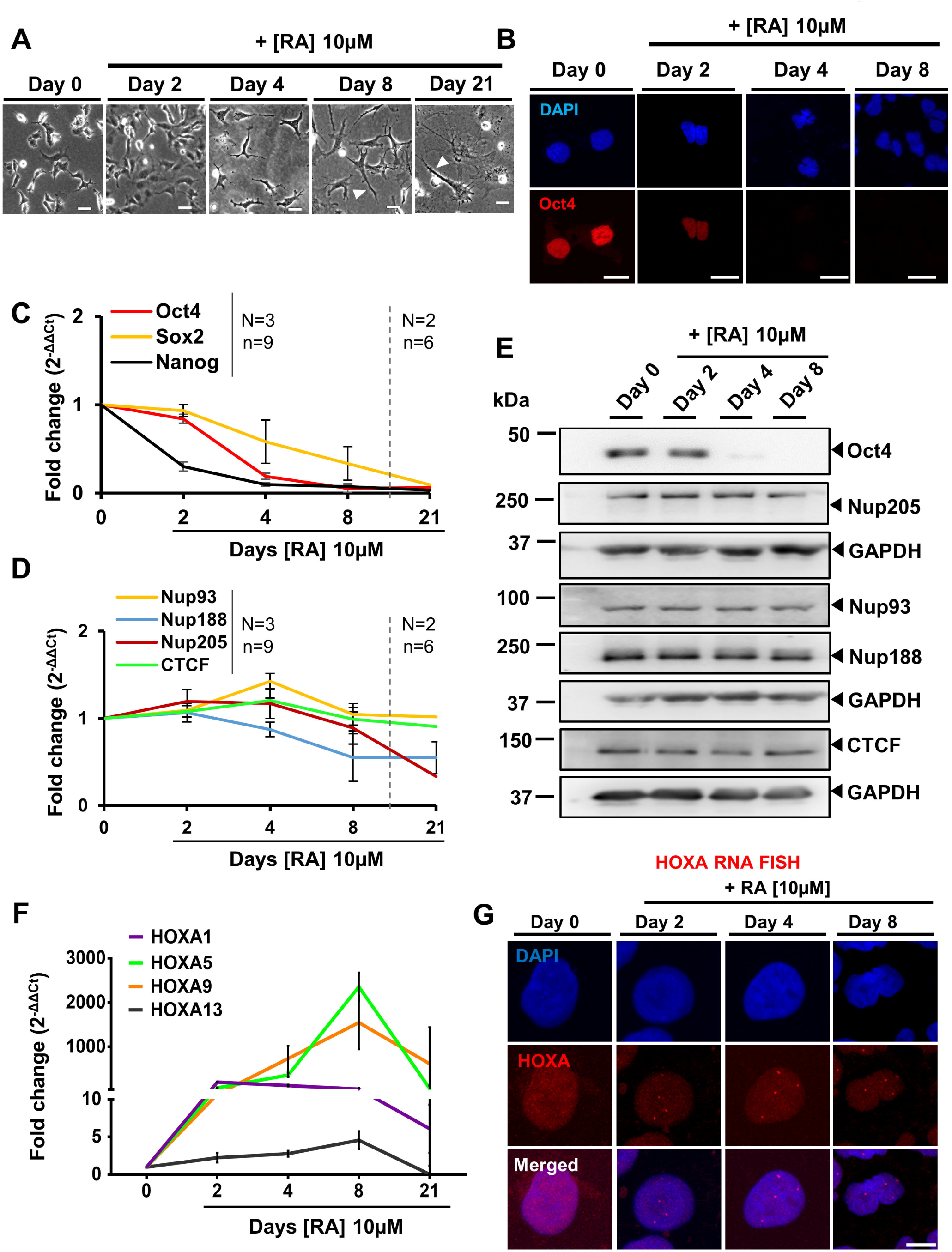
Levels of Nup93 and its interactors are maintained during differentiation. (A) Representative bright field images of NT2/D1 cells on Day 0, 2, 4, 8 & 21 during differentiation. NT2/D1 cells show altered morphology during differentiation. Scale bar = 10μm. (B) *Immunofluorescence* (IF) staining of Oct4 in NT2/D1 cells upon RA treatment (10 μM) showing a decrease in Oct4 levels after Day 2. Representative images showing nuclei stained with DAPI (blue) and Oct4 (Red). Scale bar =10μm. N=1. (C) qRT-PCR analyses showing expression levels of pluripotency markers Oct4, Sox2 and Nanog at Day 0, 2, 4, 8, and 21 upon RA treatment. The graph represents fold change (2^-ΔΔCt^) in mRNA levels normalized to control cells (siLacZ). N=3 independent biological replicates for Day 0, 2, 4 and 8, N=2 for Day 21, n=3 technical replicates of each biological replicate. Internal control: Actin, Error bars: SEM. (D) Transcript levels of Nup93, Nup188, Nup205, and CTCF do not show significant changes in their levels at Day 0, 2, 4, 8, and 21 upon RA treatment. Y-axis: Fold change (2^-ΔΔCt^) in mRNA levels normalized to control cells (siLacZ). N=3 independent biological replicates for Day 0, 2, 4 and 8, N=2 for Day 21, n=3 technical replicates of each biological replicate. Internal control: Actin, Error bars: SEM. (E) A representative western blot showing protein levels of Oct4, Nup93, Nup188, Nup205, and CTCF during RA mediated differentiation. Loading control: GAPDH. Protein levels of Nup93, Nup188, Nup205, and CTCF do not show significant variations across time. Data from N=3 independent biological replicates, (see Supplemental Fig. S3D for western blot quantification). (F) qRT-PCR analyses of representative genes from the HOXA locus during RA mediated differentiation of NT2/D1 cells. Graphs representing expression levels of HOXA1, HOXA5, HOXA9 and, HOXA13 at Day 0, 2, 4, 8 and 21 of RA treatment. The graph represents fold change (2^-ΔΔCt^) in mRNA levels normalized to Day 0. Actin was used as an internal control for normalization. Error bars: SEM, data from N=3 independent biological replicates that include n=3 technical replicate each, N=2 independent biological replicates for Day 21. (G) Representative images (maximum intensity projection of a confocal image stack) of HOXA RNA-FISH showing activation of HOXA transcripts upon RA mediated differentiation from Day 2 to Day 8. The nucleus stained with DAPI (Blue) and HOXA RNA FISH foci (Red). Scale bar = 10μm. Data from N=2 independent biological replicates.

### Temporal activation of HOXA gene expression by Retinoic Acid (RA)

HOXA gene expression is induced in a temporal manner in response to Retinoic Acid (RA) treatment in human embryonal carcinoma - NT2/D1 cells (Xu et al. 2014). We examined expression levels of HOXA genes in NT2/D1 cells over 21 days. We performed qRT-PCR for HOXA genes representing the 3’ end (HOXA1 and HOXA5) and 5’ end (HOXA9 and HOXA13), of the HOXA gene cluster (Fig. 2F). Untreated cells hardly showed any HOXA gene expression (Day 0, Fig. 2F). Interestingly, HOXA genes (HOXA1, A5, A9, and A13) were activated in a temporal manner upon RA treatment as revealed by a significant increase in their expression levels with time (Fig. 2F).

We independently examined HOXA expression by RNA-FISH, which showed (i) transcriptional activation on Day 2 of RA treatment (ii) stochastic activation of 1-4 copies of HOXA loci and (iii) predominance of 4 active HOXA RNA foci by Day 8 of differentiation at the single cell level (Fig. 2G, Supplemental Fig. S3E). This is consistent with 4 copies of HOXA gene locus in NT2/D1 cells as revealed by DNA-FISH on metaphase spreads (Supplemental Fig. S4A-E). Taken together, these results underscore the temporal activation of HOXA gene expression upon induction of cellular differentiation.

### Nup93 and CTCF depletion show antagonistic effects on HOXA gene expression

Since Nup93 sub-complex and CTCF were unaltered during differentiation, we asked if Nup93 or CTCF depletion impinges on expression levels of HOXA genes. We determined levels of HOXA genes upon siRNA mediated knockdown of Nup93, CTCF and their combined knockdowns in NT2/D1 cells (Fig. 3A; Supplemental Fig. S5A). While the entire HOXA gene cluster is upregulated in differentiated cells, upon Nup93 Kd (Labade et al. 2016), HOXA genes were differentially regulated. In undifferentiated NT2/D1 cells, 3’ HOXA genes (HOXA1 and HOXA5) were significantly upregulated, while the 5’ HOXA genes (HOXA9 and HOXA13), were downregulated (Fig. 3A). In marked contrast, CTCF knockdown showed the converse, i.e., a significant decrease in expression levels of 3’ HOXA genes (HOXA1 and HOXA5) and an increase in 5’ HOXA genes (HOXA9 and HOXA13) (Fig. 3A). Of note, the combined depletion of Nup93 and CTCF, rescued expression levels of the HOXA genes to their basal levels (Fig. 3A), further underscoring a co-regulatory role of Nup93 and CTCF in regulating HOXA expression in undifferentiated cells. Notably, transcript levels of pluripotency markers (Oct4, Sox2 and, Nanog) were unaffected upon Nup93, CTCF and their co-depletion (Fig. 3B and C). Therefore, the loss of Nup93, CTCF or both, does not alter the undifferentiated state of NT2/D1 cells. Also, Nup93 depletion did not significantly alter cell cycle profiles (Supplemental Fig. S5B). Furthermore, nucleoporins such as Nup93 and Nup153 regulate gene expression but do not drastically perturb nuclear transport (Jacinto et al. 2015; Ibarra et al. 2016; Labade et al. 2016). We, therefore, performed Nup93 knockdown in undifferentiated NT2/D1 cells and tested for effects on nuclear transport. We detected a marginal change in mRNA export upon Nup93 depletion (Supplemental Fig. S5C and D). However, Nup98 (a known mRNA export regulator), significantly inhibited mRNA export in NT2/D1 cells (Supplemental Fig. S5C and D) (Oka et al. 2010). Taken together these results reveal an antagonistic role of Nup93 and CTCF in regulating 3’ and 5’ regions of the HOXA gene locus in undifferentiated cells.

**Figure 3.**
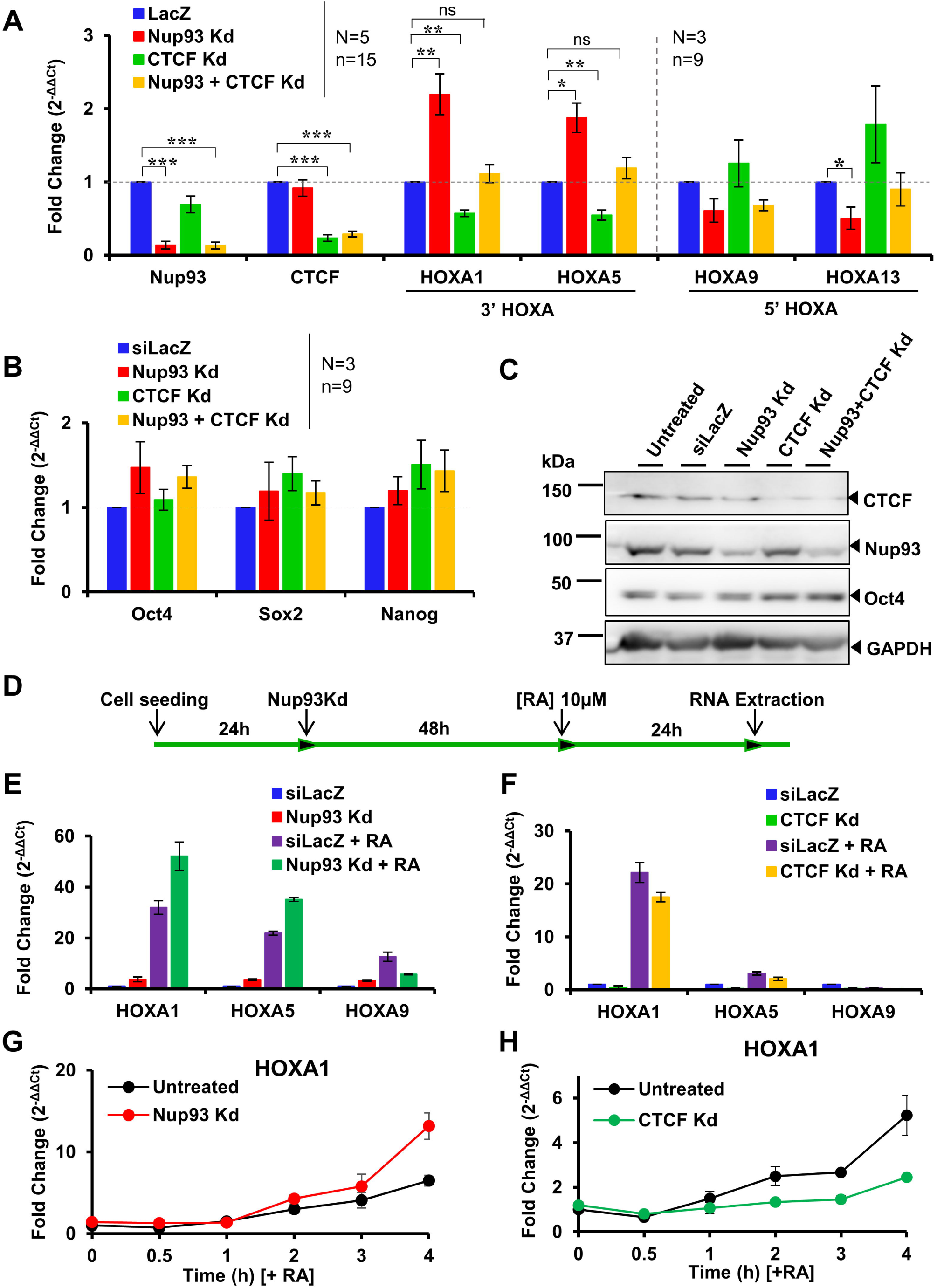
Nup93 and CTCF regulate HOXA gene expression in an antagonistic manner in undifferentiated NT2/D1 cells. (A) qRT-PCR analyses of representative HOXA genes from 3’ region (HOXA1 and HOXA5) and 5’ region (HOXA9 and HOXA13). Nup93 Kd shows a significant increase in transcript levels of 3’ HOXA genes (HOXA1 and HOXA5) and decrease in 5’ HOXA genes (HOXA9 and HOXA13). CTCF Kd shows a decrease in 3’ HOXA genes (HOXA1 and HOXA5) and increase in 5’ HOXA genes (HOXA9 and HOXA13). Graph showing, Y-axis: fold change (2^-ΔΔCt^) in mRNA levels normalized to siLacZ treated cells. Data compiled from N=5 independent biological replicates for Nup93, CTCF, HOXA1 and HOXA5, N=3 for HOXA9 and HOXA13, n=3 technical replicates in each biological replicate. Error bars: SEM. Student’s t-test (* = p<0.05, **= p<0.01, ***= p<0.001). (B) Transcript levels as determined by qRT-PCR of pluripotency genes - Oct4, Sox2, and Nanog are unaltered after 48 hours of Nup93, CTCF and double knockdowns of Nup93 and CTCF. Data compiled from N=3 independent biological replicates, n=3 technical replicates in each biological replicate, Error bars: SEM. (C) Representative western blots for CTCF, Nup93, and Oct4, showing that the relative levels of these proteins are unaffected upon Nup93, CTCF Kd or both. Loading control: GAPDH. Data compiled from N=2 independent biological replicates. (See Supplemental Fig. S5A for quantification of western blot). (D) Schematic for Nup93 knockdown followed by RA treatment of NT2/D1 cells. (E, F) Nup93 or CTCF Kd affect RA mediated induction of HOXA gene expression. A graph representing fold change (2^-ΔΔCt^) in transcript levels of HOXA1, HOXA5 and HOXA9 upon (E) Nup93 Kd and (F) CTCF knockdown followed by RA treatment. Data from N=1, with n=3 technical replicates of each biological replicate. *The second biological replicate for the same experiment is plotted independently (Supplemental Fig. S5E and F)*. (G, H) Effect of Nup93 (G) and CTCF Kd (H) on HOXA1 expression at early time points upon RA treatment. A graph representing fold change (2^-ΔΔCt^) in transcript levels (Y-axis) of HOXA1 from 30min to 4h of RA treatment (X-axis). *The second biological replicate for the same experiment is plotted independently (Supplemental Fig. S5G and H)*.

### Nup93 or CTCF depletion modulates RA induced HOXA expression

While, Nup93 and CTCF show antagonistic roles in modulating HOXA gene expression levels in NT2/D1 cells, we asked if Nup93 or CTCF depletion impinge on HOXA1 expression levels? We, therefore, performed Nup93 and CTCF knockdowns followed by RA treatment of NT2/D1 cells for 24 hours (Fig. 3D). RA treatment in Nup93 depleted NT2/D1 cells results in the enhanced activation of 3’ HOXA genes - HOXA1 and HOXA5 respectively, whereas the 5’ HOXA gene - HOXA9, was comparatively attenuated (Fig. 3E; Supplemental Fig. S5E). This suggests that the depletion of Nup93 potentially primes the 3’ end of HOXA genes for activation, that further enhances HOXA1 and HOXA5 expression upon RA treatment (Fig. 3E; Supplemental Fig. S5E). In contrast, CTCF depletion attenuates expression of 3’ HOXA genes (HOXA1 and HOXA5) in response to RA treatment (Fig. 3F; Supplemental Fig. S5F). Next, we determined if Nup93 and CTCF depletion activates HOXA1 expression at early time points (0-4h) upon RA induction? (Fig. 3G, H; Supplemental Fig. S5G, H). Interestingly, Nup93 depleted cells showed an early induction of HOXA1 (~1.5h), while in contrast, CTCF depleted cells showed a relatively delayed induction of HOXA1 (~3h) (Fig. 3G, H; Supplemental Fig. S5G, H). Of note, differentiation of ES cells shows significant variation in gene expression levels across biological replicates as reported previously (Willems et al. 2008; Mason et al. 2014), we have therefore plotted biological replicates of RA induced HOX expression upon Nup93 or CTCF depletion independently (Fig. 3E-H and Supplemental Fig. S5E-H). Taken together, these results suggest that Nup93 and CTCF depleted cells differentially respond to RA treatment even during early induction of HOXA1 expression. In summary, these results underscore opposing roles of Nup93 and CTCF on HOXA gene regulation during differentiation.

### HOXA gene locus relocates away from the nuclear periphery upon Nup93 but not CTCF depletion

The spatial localization of the HOXA gene locus correlates with its expression status (Morey et al. 2007). Since Nup93 and CTCF depletion showed antagonistic effects on HOXA expression levels, we therefore, determined the effect of Nup93 and CTCF depletion on the spatial organization of the HOXA gene locus by 3D-FISH (3Dimensional Fluorescence In-situ hybridization) (Fig. 4A). Interestingly, Nup93 knockdown showed a significant movement of the HOXA gene locus away from the nuclear periphery (Median (M) = 1.26 μm, siLacZ= 1.04 μm) (Fig. 4B). In contrast, CTCF Kd did not affect the position of the HOXA gene locus (M= 1.09 μm, siLacZ= 1.04 μm) (Fig. 4B). In addition, we also determined the effect of RA treatment on HOXA position in Nup93 and CTCF depleted cells. The HOXA gene locus showed a significant movement away from the nuclear periphery upon RA treatment consistent with its enhanced expression (Fig. 4B). Interestingly, the altered position of the HOXA gene locus upon Nup93 knockdown (M= 1.26 μm, siLacZ= 1.04 μm), was maintained even upon RA treatment (M= 1.33 μm, siLacZ= 1.4 μm) (Fig. 4B). This suggests that Nup93 depletion alone is sufficient for untethering the HOXA locus from the nuclear periphery irrespective of the induction of HOXA expression. In contrast, RA treatment in CTCF depleted cells (M= 1.28 μm, siLacZ= 1.4 μm) showed a significant movement of HOXA locus from the nuclear periphery as compared to CTCF Kd alone (M= 1.09 μm, siLacZ= 1.04 μm) (Fig. 4B). Taken together, these results underscore an overarching role of Nup93 in tethering and repressing the HOXA gene locus at the nuclear periphery, consistent with its role in differentiated cells (Labade et al. 2016).

**Figure 4.**
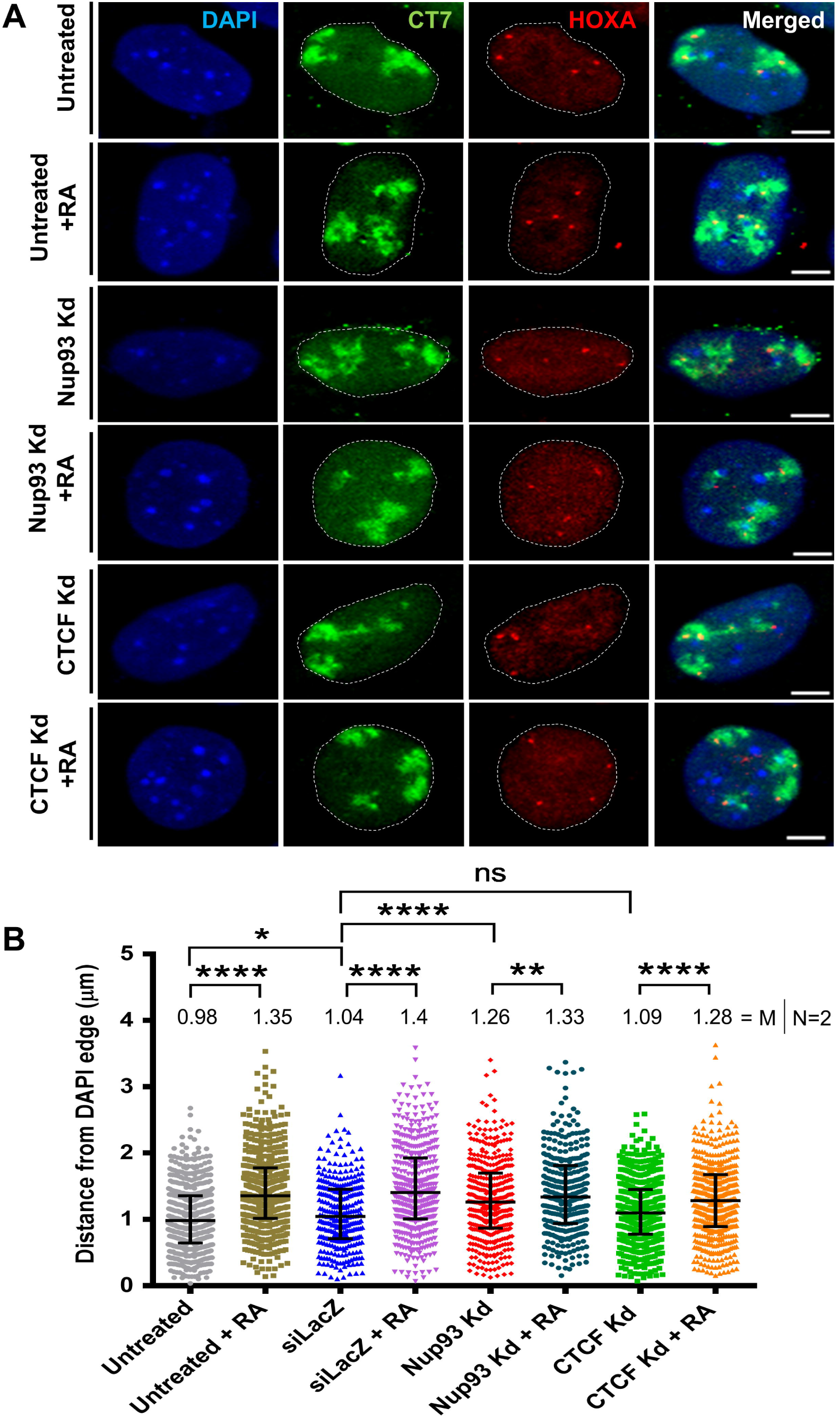
HOXA gene locus disengages from the nuclear periphery upon induction of HOXA expression. (A) Representative images of maximum intensity projections from 3D confocal image stacks of 3-Dimensional Fluorescence *In-situ* Hybridization (3D-FISH), showing nucleus stained with DAPI (blue), Chromosome Territory 7 (CT7, green) and HOXA gene locus (Red) in control and knockdown NT2/D1 cells +/-RA. Scale bar = 10μm. The nuclear boundary is represented by a white dotted line in green and red channels. The relative distance of the HOXA gene locus was measured from the nuclear border demarcated by DAPI edge, in control, RA treated, Nup93 Kd, and CTCF Kd cells. (B) Quantification of 3D-FISH data from (A). RA mediated HOXA induction shows a significant disengagement of HOXA gene locus from nuclear periphery, also seen upon Nup93 Kd, Nup93 Kd + RA, but not upon CTCF Kd. Scatter plot showing shortest distance of HOXA gene loci from edge of nucleus, demarcated by DAPI. Untreated (n=586), Untreated + RA (n=556), siLacZ (n=316), siLacZ + RA (n=449), Nup93 Kd (n=421), Nup93 Kd + RA (n=341), CTCF Kd (n=559), CTCF Kd + RA (n=485). Data from N=2 independent biological replicates, *horizontal bar* represents median with interquartile range, n: number of gene loci. Values above each scatter represent median (M). Statistical analyses: Significance calculated using Mann-Whitney U test (* = p<0.05, **= p<0.01, ***= p<0.001, ****=p< 0.0001).

### Nup93 and CTCF re-occupy HOXA gene locus upon differentiation

In order to address the role of Nup93 on HOXA1 regulation during differentiation, we ascertained if the Nup93 sub-complex (Nup188-Nup93-Nup205), maintains its stability during differentiation. We performed co-immunoprecipitation of Nup188 (interactor of Nup93) on Day 0, 4, 8 and 21 upon RA treatment. Interestingly, co-immunoprecipitation revealed that Nup188 indeed interacts with Nup93. The association between Nup93 and Nup188 was maintained throughout differentiation (Fig. 5A). However, Nup188 did not show an association with Nup205, comparable to its status in differentiated cells consistent with previous studies (Fig. 5A) (Miller et al. 2000; Theerthagiri et al. 2010; Braun et al. 2016; Labade et al. 2016). Of note, Nup188 did not show an association with CTCF (established regulator of HOXA). In summary, the stability of the Nup93 sub-complex was maintained during differentiation.

**Figure 5.**
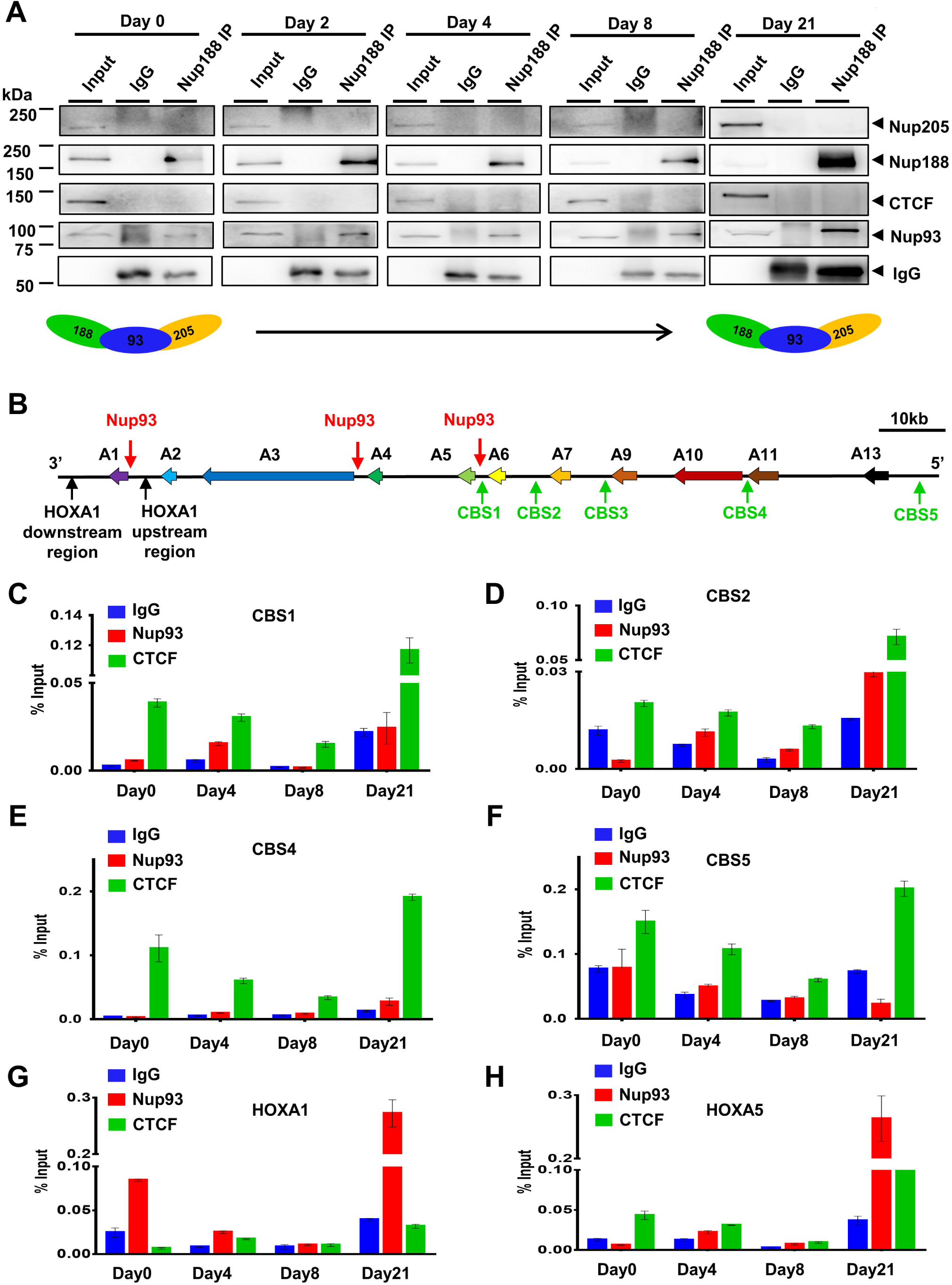
Occupancy of Nup93 and CTCF on HOXA locus during differentiation. (A) Co-immunoprecipitation of Nup188 during differentiation at Day 0, 2, 4, 8 and 21. Nup188 interacts with Nup93 but not with Nup205 or CTCF (Data from a single experiment, N=1). Model representing stable Nup93 sub-complex from Day 0 to Day 21. (B) Schematic representation of HOXA gene locus (HOXA1-HOXA13) showing location of Nup93 (Red arrows) and CTCF binding sites (CBS, blue arrows) on 3’-end and 5’-ends of HOXA gene locus. (C-F) CTCF ChIP-qPCR shows gradual decrease in CTCF occupancy on its conserved binding sites (C) CBS1, (D) CBS2, (E) CBS4 and (F) CBS5 upon RA treatment on Day 0, 4 and 8. CTCF showed an increase in its occupancy on all CBSs on Day 21. Y-axis: immunoprecipitated DNA relative to 1% input (N=1, data from one experiment that include a total of three technical replicates), error bar = standard error of the mean (SEM). (*Biological replicate 2 is plotted separately in Supplemental Fig. S6)*. (G, H) Nup93 ChIP-qPCR shows gradual decrease in its occupancy on (G) HOXA1 promoter and (H) HOXA5 promoter on Day 0, 4 and 8 during differentiation of NT2/D1 cells in response to RA treatment. Nup93 re-occupies 3’ HOXA genes (HOXA1 and HOXA5) upon differentiation of NT2/D1 cells at Day 21. Y-axis: immunoprecipitated DNA relative to 1% input (N=1, data from one experiment that include a total of three technical replicates), Error bar = standard error of the mean (SEM). (*Biological replicate 2 is plotted separately in Supplemental Fig. S6)*.

Nup93 and CTCF have specific binding sites on the HOXA locus (Fig. 5B) (Labade et al. 2016). Nup93 binding sites are predominantly enriched on 3’ HOXA genes (HOXA1, HOXA3, and HOXA5), while CTCF binding sites are enriched on 5’ HOXA genes (Fig. 5B). We determined the association of Nup93 and CTCF with HOXA gene locus during differentiation. We performed Nup93 and CTCF ChIP on Days 0, 4, 8 and 21 of differentiation and determined the relative enrichment of CTCF and Nup93 on CTCF binding sites (CBSs) and 3’-end HOXA genes (HOXA1 and HOXA5) respectively by ChIP-qPCR (Fig. 5C-H). Interestingly, CTCF showed a gradual decrease in its occupancy on all CTCF binding sites (CBSs) upon RA treatment until Day 8 (Fig. 5C-F; Supplemental Fig. S6A-D). RA prevents CTCF binding on HOXA, thereby promoting HOXA expression specifically during differentiation (Oh et al. 2018). Remarkably, CTCF showed a striking increase in its occupancy on CTCF binding sites on Day 21 (Fig. 5C-F). We did not observe an enrichment of Nup93 on the CTCF binding sites (except CBS2 on Day 21) on Days 0, 4, 8 and 21, suggesting reduced occupancy of Nup93 at the 5’-end of the HOXA gene cluster (Fig. 5C-F). Nup93 is enriched on HOXA1 and HOXA5 promoters on Day 0, with a decrease in its occupancy upon differentiation (Day 4 and Day 8) (Fig. 5G and H; Supplemental Fig. S6E and F). Remarkably, Nup93 showed a re-enrichment on HOXA1 promoter on Day 21 of differentiation (Fig. 5G and H). This suggests that Nup93 associates with and represses HOXA gene locus on Day 0 (undifferentiated) and Day 21 (differentiated) respectively. To ascertain the binding specificity of Nup93 and CTCF, we examined their occupancy on regions outside the HOXA1 promoter (~2 kb upstream and downstream of the HOXA1 promoter region) (Fig. 5B; Supplemental Fig. S7A and B; Supplemental Fig. S8A and B). We found that both Nup93 and CTCF do not associate with sites immediately upstream of the HOXA1 promoter (~2 kb) (Supplemental Fig. S7A and B; Supplemental Fig. S8A and B). A control ChIP with anti-PanH3 antibody (anti-core histone H3), revealed an enrichment on HOXA1 and HOXA5 promoters and CTCF binding sites, reflecting the pull-down efficiency of the ChIP experiment (Supplemental Fig. S7C-J; Supplemental Fig. S8C-J). Taken together, these findings suggest co-regulatory roles of Nup93 and CTCF in the organization of HOXA locus during differentiation.

### HOXA gene locus re-localizes closer to the nuclear periphery upon differentiation

The HOXA gene locus is reorganized during transcriptional activation (Ferraiuolo et al. 2010; Rousseau et al. 2014). We showed that the untethering of the HOXA gene locus from the nuclear periphery correlates with its upregulation upon Nup93 Kd in differentiated cells (Labade et al. 2016). Since HOXA genes were expressed upon RA treatment, we visualized the spatial localization of the HOXA gene locus during differentiation using 3D-FISH (Fluorescent In situ hybridization) (Fig. 6A and B). The fluorescently labelled HOXA gene locus is proximal to the nuclear periphery on Day 0 (M= 1.12 μm) (Fig. 6B and C). Interestingly, HOXA gene locus moved significantly away from the nuclear periphery on Day 2 (M= 1.61 μm) and Day 4 (M=1.5 μm) of differentiation respectively (Fig. 6B and C). Remarkably, HOXA gene locus showed a significant relocation proximal to the nuclear periphery on Day 8 (M= 1.18 μm) and Day 21 (M= 1.13 μm), consistent with its re-repression on Day 21 of differentiation (Fig. 6B and C). Taken together, these results suggest that the activation and repression of HOXA gene locus during differentiation is associated with its untethering and re-tethering with the nuclear envelope.

**Figure 6.**
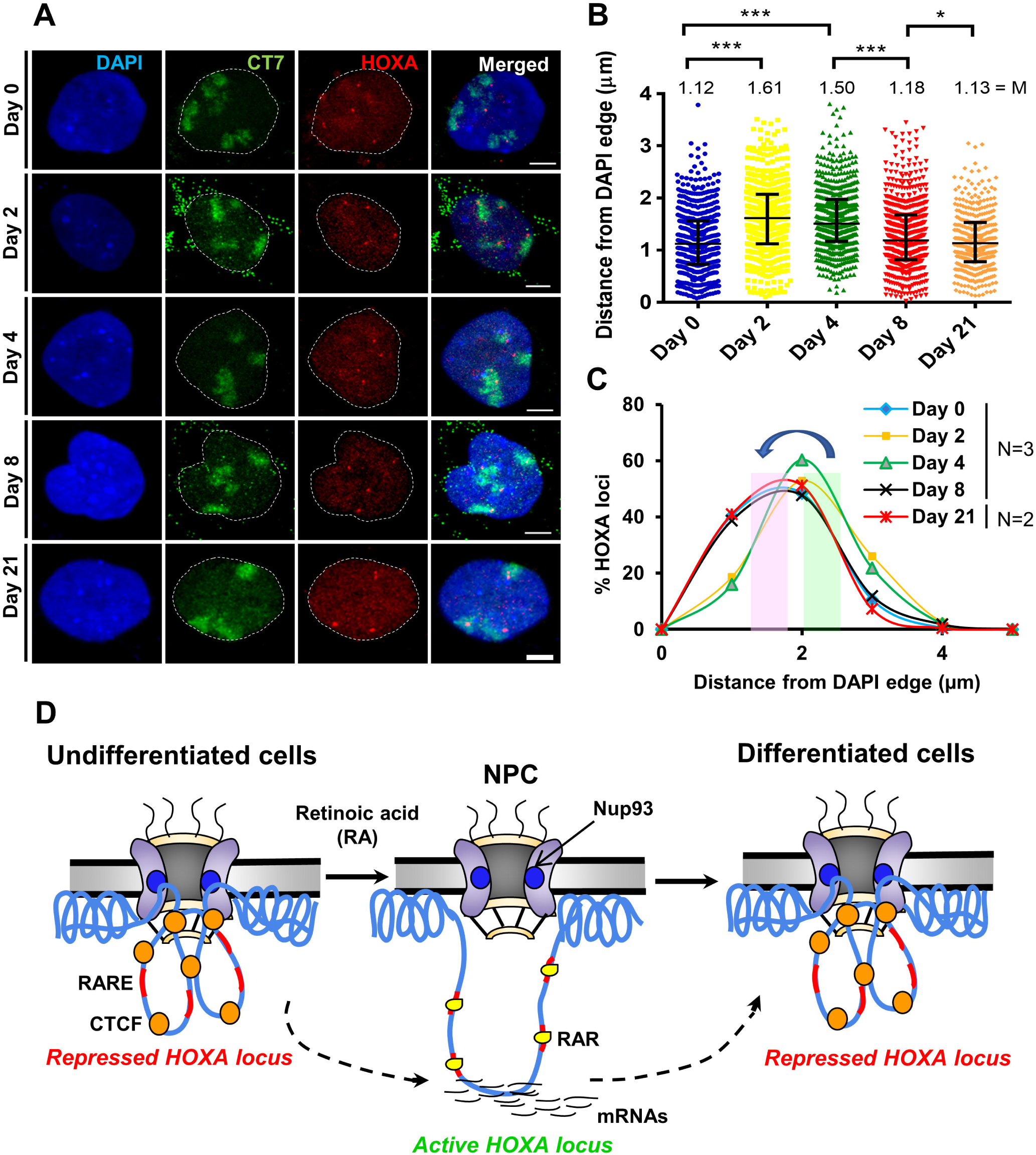
HOXA gene loci are restored closer to the nuclear periphery upon differentiation. (A) Representative images showing maximum intensity projections of 3D confocal image stacks with the nucleus labelled with DAPI (blue), Chromosome Territory 7 (CT7, green) and HOXA gene locus (Red) in NT2/D1 cells treated with RA and images acquired at specific time points (Days 0, 2, 4, 8, and 21) of RA treatment. (B) Scatter plot showing shortest distance (μm) of HOXA gene locus from DAPI edge, across days as indicated on the Y-axis. Data compiled from N=3 biological replicates for Day 0 (n=537), Day 2 (n=548), Day 4 (n=601) & Day 8 (n=621) and N=2 for Day 21 (n=400), n: number of gene loci. *Horizontal bar* represents the median with interquartile range. Values above each scatter represent the median (M). Statistical significance was calculated using Mann-Whitney U test (* = p<0.05, **= p<0.01, ***= p<0.001). (C) Data from (B) plotted as % frequency distribution of the spatial location of HOXA loci from the edge of the nucleus across days upon RA treatment in bins of 1μm. Y-axis: % HOXA loci. HOXA gene loci show a distinctive disengagement from the nuclear periphery upon RA mediated differentiation (Day 2 and Day 4). Remarkably, HOXA gene locus re-localizes closer to the nuclear periphery from Day 8 and Day 21 of differentiation. The nuclear boundary is represented by a white dotted line, Scale bar = 10μm. (D) Speculative model of the spatiotemporal organization and function of the HOXA gene cluster during differentiation. In undifferentiated NT2/D1 cells, HOXA gene cluster is positioned proximal to the nuclear periphery, potentially sequestered by Nup93 and CTCF, maintained in a relatively repressed state. Retinoic acid-mediated induction of HOXA expression, is accompanied by a movement of HOXA gene loci away from the nuclear periphery. Retinoic acid (RA) induces Retinoic acid receptor (RAR) which further associates with Retinoic acid response elements (RARE) that evicts CTCF from CTCF binding sites (CBSs). This disrupts chromatin loops within the HOXA gene locus. Upon differentiation, HOXA gene locus relocates closer to the nuclear periphery and is re-occupied by Nup93 and CTCF, consistent with its re-repression.

## DISCUSSION

The central role of nucleoporins is to regulate the transport of RNA and proteins. However, nucleoporins are also involved in gene regulation. Nucleoporins - Nup93, Nup153, Nup98, Nup50, Seh1 and Nup210 (Buchwalter et al. 2014; Pascual-Garcia et al. 2017; Liu et al. 2019), directly associate with developmental genes and regulate developmental processes (Liang et al. 2013; Jacinto et al. 2015; Ibarra et al. 2016). This implicates NPCs potentially as a stable anchor for developmental genes among others, in the nucleus. However, the physiological relevance of nucleoporin-chromatin associations in modulating the organization and activity of genes during differentiation is unclear. Here we show that nucleoporin Nup93 modulates the organization and function of the HOXA gene locus during differentiation. ChIP-seq analysis of Nup93 - a core component of the nuclear pore complex reveals that Nup93 shares binding sites with the genome organizer - CTCF, suggesting a functional overlap between Nup93 and CTCF in the organization and regulation of the HOXA gene cluster during differentiation.

Nup93 is a stable nucleoporin with relatively low dissociation rates from the nuclear pore complex (Rabut et al. 2004). ChIP-seq analysis revealed a genome-wide occupancy of Nup93 in differentiated cells (Fig. 1; Supplemental Fig. S1). Notwithstanding its localization at the nuclear membrane, we did not find a correlation between Nup93 occupancy with gene-poor chromosomes, localized toward the nuclear periphery, considering that gene poor chromosome territories are closer to the nuclear envelope in the interphase nucleus (Supplemental Fig. S1E and F). Surprisingly Nup93 hardly showed any occupancy on human Chr.X, despite its relatively peripheral nuclear localization (Supplemental Fig. S1E and F) (Chen et al. 2016). This is consistent with Dam-ID analyses of Nup93 from U2OS cells, which does not reveal Nup93 occupancy on X-chromosomes (Ibarra et al. 2016). We surmise that an active mechanism excludes Nup93 occupancy from human X-chromosome.

In addition to Nup93, other nucleoporins such as Nup153, Nup50, Nup210, Seh1, and Nup98 are implicated in the regulation of developmental gene expression (Kalverda et al. 2010; Liang et al. 2013; Buchwalter et al. 2014; Jacinto et al. 2015; Liu et al. 2019). It is interesting that Nup93 associated genes are enriched for developmental pathways such as Hippo signaling and Wnt-signaling pathway (Supplemental Fig. S2A). Moreover, Nup93 binding shares consensus DNA binding motifs with that of transcription factors such as - EGR3, ZN394, ZN502, MAZ, RREB1, and FOXG1, also associated with development (Supplemental Fig. 1C; Supplemental Fig. S2B). In summary, Nup93 occupancy overlaps with transcription factor binding sites associated with the regulation of developmental genes.

Furthermore, ChIP-seq analyses of Nup93 revealed its overlap with CTCF (Fig. 1E-G; Supplemental Fig. S2C and D), consistent with CTCF binding motif enriched on Nup93 binding sites (Brown et al. 2008). These results suggest the intriguing possibility that Nup93 potentially modulates topological chromatin contacts such as insulator or enhancer-promoter contacts mediated by CTCF (Handoko et al. 2011; Narendra et al. 2015; Oh et al. 2018). Interestingly, Nup2 and Nup98 functions as insulators in *S.cerevisiae* and *Drosophila*, similar to CTCF (Ishii et al. 2002; Kalverda and Fornerod 2010). Nup98 - although an off-pore nucleoporin, promotes enhancer-promoter contacts along with CTCF, as shown in *Drosophila* S2 cells (Pascual-Garcia et al. 2017). Furthermore, genome-wide analysis of Nup93 binding revealed that Nup93 associates with enhancer marks such as H3K4me1 and H3K27ac (Fig. 1D and E) (Creyghton et al. 2010). H3K27ac mark is associated with Nup93, as revealed by DamID-Seq analyses (Ibarra et al. 2016). These findings further implicate Nup93 as a stable tether that facilitates long-range chromatin contacts between enhancers and promoters of genes involved in development. Considering the major role of CTCF in regulating cell type-specific enhancer-promoter contacts (Ren et al. 2017), we surmise that Nup93 potentially functions as an anchorage site for nucleating proteins that further promote cell type and cell state specific chromatin organization.

Our studies reveal a remarkable molecular interplay between Nup93 and CTCF in the organization of the HOXA gene locus during differentiation. The depletion of CTCF in undifferentiated NT2/D1 cells downregulates 3’ HOXA gene expression (HOXA1 and HOXA5) (Fig. 3A). However, CTCF depletion in differentiated cells does not largely perturb the HOXA gene cluster but selectively overexpresses HOXA13 (Labade et al. 2016). In contrast, Nup93 depletion upregulates HOXA genes, suggesting a predominant role of Nup93 in regulating the organization of the HOXA gene cluster, in a CTCF independent manner in differentiated cells. Nup93 and CTCF exhibit a more proactive role in undifferentiated cells, as they show antagonistic functions in regulating the expression of 3’ and 5’ ends of HOXA gene locus (Fig. 3A, E and F). Interestingly, CTCF depletion in monocytes from peripheral blood (THP-1 cells), showed a decrease in HOXA expression levels (Crutchley 2014). Similarly, CTCF depletion showed decreased expression of homeotic genes during early stages of *Drosophila* development (Mohan et al. 2007). In contrast, CTCF depletion upregulates 5’ but not 3’ end of the HOXA gene cluster in NT2/D1 cells (Xu et al. 2014). Interestingly, 3’ HOXA genes are relatively more responsive to retinoic acid (RA) treatment in a Nup93 depleted background (Fig. 3E). We surmise that Nup93 depletion enhances the accessibility of the Retinoic Acid Response Element (RARE) within the HOXA gene cluster, to the retinoic acid receptor (RAR) thereby making HOXA more responsive to RA treatment (Fig. 6D). We found that HOXA induction either by RA treatment or Nup93 but not CTCF depletion untethers the HOXA gene locus from the nuclear periphery (Fig. 4A and B). We speculate that the depletion of CTCF alone is insufficient to displace the HOXA gene locus from the nuclear periphery in the presence of Nup93.

The temporal and collinear expression of HOXA genes is essential for normal development and differentiation (Narendra et al. 2015; Wang et al. 2015; Oh et al. 2018). CTCF functions as an organizer of chromatin loops of the HOXA gene cluster which facilitates the spatiotemporal control of HOXA gene expression during differentiation (Narendra et al. 2015; Wang et al. 2015; Hansen et al. 2017; Oh et al. 2018). However, the mechanism underlying the regulated control of HOXA gene expression during differentiation is unclear. Notably, both Nup93 and CTCF co-occupy the HOXA gene locus, suggesting an uncharacterized mechanism involving Nup93 in the spatiotemporal regulation of HOXA gene expression. Consistent with the occupancy profiles of Nup93 on HOXA locus, Nup93 represses HOXA genes in undifferentiated and differentiated cells (Fig. 6D). We found that CTCF is enriched on the HOXA gene locus in undifferentiated NT2/D1 cells (Fig. 5C-F). However, upon induction of HOXA expression, CTCF occupancy on CTCF binding sites (CBSs) decreases, accompanied by HOXA expression, limited to the specific time window of differentiation. Consequently, HOXA expression is attenuated upon differentiation and in differentiated cells, potentially mediated by the repressive effects and re-tethering of HOXA by Nup93 (Fig. 6D).

It is remarkable that the HOXA gene locus shows a distinct untethering from the nuclear envelope during differentiation (Fig. 6A and B). Untethering of HOXA locus may facilitate the dynamic reorganization of chromatin domains within HOXA locus upon activation as revealed by chromosome conformation (3C) studies (Ferraiuolo et al. 2010; Rousseau et al. 2014; Xu et al. 2014). The HOXA gene cluster is in a relatively poised state and is rapidly activated when it receives developmental cues such as RA (Xu et al. 2014). This is consistent with altered interchromatin contacts of the HOXA gene cluster during differentiation of THP1 monocytes (Rousseau et al. 2014). We speculate that the HOXA gene locus transitions from a repressive chromatin compartment (nuclear periphery) to a relatively more active compartment (away from the nuclear periphery) during differentiation accompanied by the activation of HOXA gene locus (Fig. 6D) (Narendra et al. 2015). In summary, these studies suggest a remarkable coordination between Nup93 and CTCF in the spatiotemporal organization and function of the HOXA gene locus during differentiation.

In conclusion, this study unravels a crosstalk between nucleoporin Nup93 and the genome organizer CTCF in the regulation of HOXA expression during differentiation. We show that Nup93 associated genes are involved in development and differentiation. Nup93 bound genomic regions are enriched for enhancer marks such as H3K4me1 and H3K27ac suggesting its role in promoting enhancer-promoter contacts. Interestingly, Nup93 occupancy strongly correlates with the genome organizer-CTCF, underscoring its role in genome organization. Furthermore, Nup93 or CTCF knockdown shows antagonistic roles in regulating 3’ and 5’ regions of the HOXA gene cluster in undifferentiated cells. Remarkably, RA mediated induction of HOXA expression shows decreased occupancy of Nup93 and CTCF on HOXA locus during differentiation and their re-enrichment in differentiated cells. This further correlates with the untethering and re-tethering of the HOXA gene locus with the nuclear periphery. Taken together, these results provide key insights into the mechanisms that regulate the spatiotemporal activation and inactivation of the HOXA gene cluster during differentiation.

## METHODS

### Cell culture

NTERA-2 cl. D1 (NT2/D1) cells were obtained from the lab of Sanjeev Galande (IISER Pune) with permission from Peter Andrews (The University of Sheffield, UK). Cells were cultured in Dulbecco’s modified Eagle’s medium (DMEM) (Gibco, 11995) supplemented with 10% fetal bovine serum (Sigma, 2959), 100 U/ml penicillin, 100 g/ml streptomycin (Gibco, 15070-063), and 2mM L-glutamine (1X Gibco^®^ GlutaMAX™ Supplement, 35050061) at 37°C with 5% CO_2_. For differentiation, NT2/D1 cells treated with 10μM retinoic acid (RA). We ensured that cultures were free of Mycoplasma contamination, by monitoring DAPI stained cells periodically. The authenticity of NT2/D1 cells was validated by karyotyping. NT2/D1 cells consistently show modal chromosome numbers of 62-63 (Supplementary Fig. S3B and C).

### Transient siRNA mediated knockdown

Transient knockdowns were performed using siRNA oligonucleotides from Dharmacon, USA. Briefly, NT2/D1 cells (~0.2 × 10^6^) were seeded in 6-well plate 24 hours prior to transfection for cells to attain confluency of ~50-60%. The cells were transfected with siRNA oligonucleotides at the final indicated concentrations (Nup93 - 20nM, Nup188 - 20nM, Nup205 - 50nM, CTCF-50nM) using RNAiMax (Invitrogen, 13778) in reduced serum Opti-MEM (Gibco, 31985). Medium containing transfection mix was replaced with complete DMEM 24h post-transfection. Cells were harvested after 48 h or 72 h of knockdown and processed for Western blots or RNA extraction. siLacZ (Dharmacon) was used as a control. siRNA sequences are given in Supplemental Table S1.

### Immunofluorescence Assay

Cells were washed with 1X PBS and fixed with 4% PFA for 10 minutes at RT. Cells were permeabilized with 0.5% Triton X-100 for 10 minutes followed by blocking with 1% BSA/ 1X PBST (1X PBS +0.1% Tween 20) for 1h at Room temperature (RT). Cells were incubated with primary antibody (Anti Oct4 antibody, DHSB, Cat: PCRP-POU5F1-1D2) diluted in 1% BSA (in 1XPBST) for 2hr at RT in a humidified chamber. After 3 washes with 1X, PBS cells were incubated with secondary antibody (Anti-mouse antibody–Alexa Fluor 568, Molecular Probes Cat: A11004) diluted in (1:1000) 1% BSA (in PBST) for 1h at RT in a humidified chamber. Cells were washed with 1X PBS and stained with DAPI (0.1 μg/ml in 2X FA/SSC) for 5 min at RT. Coverslips were mounted on glass slides using Slowfade Gold antifade. Image acquisition was performed on a Zeiss LSM710 confocal microscope with a 63X Plan-Apo1.4 NA oil immersion objective with 405-nm, 488-nm, and 561-nm laser at 1 to 2.5 digital zoom. Detailed image analysis protocol is in the Supplementary material.

### Western Blotting

Western blot was performed as described previously (Labade et al. 2016). Antibodies used for western blotting: Rabbit anti–Nup93 (1:500, sc-292099, Lot-E0211, Santa Cruz Biotechnology, CA), rabbit anti-Nup188, (1:1000, Abcam, ab86601, Lot-GR43443-4), rabbit anti-Nup205 antibody (1:500, HPA024574, Lot-R11937, ATLAS antibodies) and rabbit anti-CTCF antibody (1:500, 07-729, Lot-2375606, Millipore). Secondary antibodies: Donkey anti-rabbit immunoglobulin G horseradish peroxidase (1:10,000, GE NA9340V) and Sheep anti-mouse immunoglobulin G-HRP (1: 10,000, NA9310V) were diluted in 0.5% milk (1X TBST). Blots were developed using ECL Prime (Amersham 89168-782) at incremental exposures of 10 seconds acquired using LAS4000 (GE). Densitometry analysis of western blots was performed using ImageJ software from three independent biological replicates. GAPDH was used as an internal control for normalization. Detailed western blot protocol is in the Supplementary material.

### Co-Immunoprecipitation (Co-IP)

For Co-IP, cells were lysed in Co-IP buffer (50 mM Tris HCl, pH 8.0, 150 mM NaCl and 0.5% NP-40) supplemented with protease inhibitor cocktail. Lysates were pre-cleared using protein-A Dynabeads (Invitrogen, 10002D), 1 hour at 4°C. After pre-clearing, total protein was quantified using BCA assay. Nup188 antibody (2μg), was incubated with 15μl Protein-A Dynabeads in 500μl 1X PBST (0.1% Tween 20 in 1X PBS) for 1h at 4°C. Antibody-coated Dynabeads were incubated overnight at 4°C with 500μg of total protein in lysis buffer on end to end rotor at 9rpm. Dynabeads were washed 9 times with Co-IP lysis buffer. Samples for western was prepared in Laemmli buffer.

### Reverse transcription-PCR and real-time quantitative PCR

Cells were washed with 1X PBS and total RNA was extracted using the Trizol method (Rio et al. 2010), cDNA was synthesized from total RNA with the ImProm-II reverse transcription system. RT-PCR was carried out using intron-exon junction primers (Supplemental Table S1). β-actin was used as internal control. Real-time quantitative PCR was performed using Bio-Rad RT-PCR instrument (CFX96 Touch) in 5 μl of reaction mixture containing KAPA SYBR Green RT-PCR mix and 2 μM each of the forward and reverse primer respectively. Fold change was calculated by double normalization of Ct values to the internal control and untreated samples by 2^-ΔΔCt^ method (Livak and Schmittgen 2001). All primers sequences are given in Supplemental Table S1.

### Three-dimensional fluorescence in situ hybridization (3D-FISH)

Bacterial artificial chromosome (BAC) DNA for HOXA locus (RP11-1132K14 for HOXA1-A9 and RP11-163M21 for HOXA*3*-A13) were purchased from CHORI BACPAC Resources (Supplemental Fig. S4A). BAC DNA extraction was performed using Hi-Pure Plasmid DNA Extraction Kit (Invitrogen K210017). Nick translation of BAC DNA was performed using the Nick Translation kit (Roche 11 745 808 910). Nick-translated DNA was precipitated using ethanol precipitation and resuspended in hybridization mix (50% deionized Formamide + Master mix containing 10% Dextran Sulphate, 0.1 mg salmon Sperm DNA in 2X SSC solution, pH 7.4). Detailed 3D-FISH and RNA FISH protocol is in the Supplementary material.

### Chromatin Immunoprecipitation (ChIP)-qPCR and ChIP-sequencing

ChIP was performed as described previously (Labade et al. 2016). In brief, cells (~1.0 ×10^7^) were cross-linked using 1% formaldehyde for 10 min at RT. ChIP was performed using antibody against Nup93 (sc-292099, Santa Cruz Biotechnology) and CTCF (07-729, Millipore). Fold enrichment over input was calculated by % input method from 2^-ΔΔCt^ (Livak and Schmittgen 2001; Haring et al. 2007). All ChIP-qPCR experiments were performed at least in two independent biological replicates as recommended in the ENCODE guidelines (Landt et al. 2012). ChIP-qPCR primers used in this study are listed in Supplemental Table S1. Nup93 ChIP-DNA (Two independent biological replicates) and Input DNA was outsourced for high throughput paired end sequencing to Genotypic Technology, Bangalore, India. Detailed ChIP-qPCR protocol and ChIP-seq analysis is in the Supplementary material. Analyzed data is available in the Supplemental Table S1.

### Statistical analysis

For qRT-PCR experiments, statistical significance was calculated using Student’s t-test with one-tailed distribution and assuming unequal variance in Microsoft Excel. For DNA FISH and Poly-A FISH experiments, significance was calculated using the Mann-Whitney U test between control and treatment samples assuming unequal variance using GraphPad Prism 6 (Version 6.01).

## DATA ACCESS

ChIP-seq datasets are deposited in the Gene Expression Omnibus with accession number GSE130656. Reviewer’s link:https://www.ncbi.nlm.nih.gov/geo/query/acc.cgi?acc=GSE130656

Token: **ensnwiwmdrybjsb**

## ACKNOWLEDGMENTS

We are grateful to IISER-Pune for all facilities and support for microscopy and imaging. We thank labs at IISER Pune for their generous contribution of reagents. We thank all members of CBL, IISER-Pune for their critical comments on the manuscript.

## AUTHORS’ CONTRIBUTIONS

AL, AS and KS designed the experiments. AL and AS performed the experiments. AL performed ChIP-seq experiment and analyzed the data with help from KK under the supervision of KS. AL, AS, KS, and KK analyzed the data and wrote the manuscript. All authors read and approved the final manuscript.

## SUPPLEMENTAL INFORMATION

**Supplementary Information. pdf:** Includes Supplementary materials and methods, Supplementary figure legends and supplementary figures.

**Additional File1.xls:** Nup93 ChIP seq peaks and associated genes.

