## Supplemental File 1 for "Nup93 and CTCF co-modulate spatiotemporal dynamics and function of the HOXA gene cluster during differentiation"

#### SUPPLEMENTAL INFORMATION

- **SUPPLEMENTARY MATERIALS AND METHODS**
- **SUPPLEMENTAL TABLE S1**
- **SUPPLENTAL REFRENCES**
- **SUPPLEMENTARY FIGURE LEGENDS**
- **SUPPLEMENTAL FIGURES**

#### SUPPLEMENTARY MATERIALS AND METHODS

##### Metaphase spread preparation

Cells were grown to a confluency of ~60% and arrested at metaphase with 0.1 µg/ml Colcemid (Roche 10 295 892 001) for 90 min. Cells were harvested by trypsinization and treated with hypotonic solution (using 0.075 M KCl) for 30 min. Hypotonic treatment was terminated by fixing cells with 4–5 drops of fixative (Methanol: Acetic Acid, 3:1), followed by centrifugation at 1000 rpm for 10 min at 4°C. The cell pellet was washed three times with a fixative solution and resuspended in a fresh fixative solution. Cells were dropped from a height onto clean glass slides. Metaphases were stained with DAPI (0.05µg/ml in 1X PBS, pH 7.0).

##### Western Blotting

Cells were lysed in RIPA buffer (50mM Tris-Cl pH 7.4, 150mM NaCl, 0.1% SDS, 0.1% sodium azide, 0.5% Na-deoxycholate, 1mM EDTA, 1% NP-40, 1X Protease inhibitor cocktail) and centrifuged at 13,000 g for 10 min at 4˚C. Total protein was estimated using a BCA kit (cat no.23225). Protein samples (20 µg) were prepared in 1X Laemmli buffer (Tris-HCl pH 6.8, 2% SDS, 20% glycerol, 0.2% bromophenol blue, 0.025% β-mercaptoethanol) and denatured at 95˚C for 5 minutes. Proteins were resolved by SDS-PAGE and transferred to activated polyvinylidene fluoride membrane (PVDF, Millipore, cat no. IPVH00010), followed by blocking with 5% non-fat dried skim milk/1X TBST (Tris-buffer saline, 0.1% Tween 20) for 1 hour at RT. Primary antibodies were diluted in 0.5% milk/1X TBST buffer. All antibody dilutions are within the linear range of detection.

**Fluorescence Assisted Cell Sorting**

Cells were trypsinized and washed with 1X PBS. Fixation was performed with chilled 70% ethanol which was added to cells with tapping. Cells were centrifuged at 1000 rpm at 4˚C for 10 min. RNAse A (1µg/µl) treatment was performed for 45 min at 37$^{o}$C. Cells were filtered through a 0.45µm filter. Propidium Iodide(10µg) was added to cells and cells were sorted using FACscalibur flow cytometer. An analysis was performed using BD Cellquest Pro. Representative images were created using Flowing Software (Turku lab).

**Poly(A) Fluorescence In situ Hybridization (FISH)**

Cells (~0.2 x $10^{6}$) were seeded on coverslips in a 6-well plate. After 48 h of Nup93 knockdown, the cells were fixed with 4% PFA in 1X PBS (pH 7.4) for 15 minutes at RT. Cells were re-fixed and permeabilized with chilled methanol for 5 minutes, followed by incubation in 2X SSC at RT for 10 minutes. Cells were hybridized with 100 µl of hybridization mix (40% Formamide, 10% Dextran Sulphate, 0.1 mg salmon Sperm DNA and 5 ng/ml of FAM oligo dT prepared in 2X SSC solution) at 37°C for 3 hours. Coverslips were washed twice with 2X SSC, followed by washes with 0.1X SSC. Cells were stained with DAPI and mounted in an antifade solution. Images were acquired using confocal microscopy using a 63X objective/N.A. 1.4 using 488nm and 405nm lasers, zoom set to 1.0. The mean fluorescence intensity of the FISH signal was determined for each cell and nucleus (demarcated by DAPI) and expressed as a ratio of the nuclear to cytoplasmic fluorescence intensity. Nuclear/cytoplasmic (N/C) fluorescence intensity ratios were calculated and plotted using GraphPad Prism software. Statistical analysis was performed using the Mann-Whitney U test.

**Three-dimensional fluorescence in situ hybridization (3D-FISH)**

NT2/D1 cells (~0.2 × 106) were seeded on coverslips in a six-well plate. After 48h of Nup93 (20 nM), CTCF (50 nM) or Nup93 (10 nM) +CTCF (50nM) knockdowns, the cells were washed with ice-cold 1X PBS and treated with cytoskeletal (CSK) digestion buffer (0.1 M NaCl, 0.3 M sucrose, 3 mM MgCl2, 10 mM PIPES (pH 7.4), 0.5% Triton X-100) for 5 min followed by fixation with 4% PFA in 1X PBS (pH 7.4) for 10 min at RT. The cells were permeabilized in 0.5% Triton X-100 (prepared in 1X PBS) for 10 min and incubated in 20% glycerol (prepared in 1X PBS) for 60 min followed by four freeze-thaw cycles in liquid nitrogen. The cells were washed three times with 1X PBS and treated with 0.1 N HCl for 10 min followed by three washes in 1X PBS for 5 min each. The cells were incubated in 50% formamide (FA)/2X saline sodium citrate (SSC) (pH 7.4) overnight at 4°C or until used for hybridization. Cells were hybridized with 3 µl of human whole chromosome 7 paint (Applied Spectral Imaging (ASI), Israel, or MetaSystems, USA) and nick-translated BAC DNA probe for HOXA gene locus (3 µl). Post-hybridization, coverslips were washed in 50% FA/2X SSC (pH 7.4), thrice for 5 min each at 45°C, followed by three washes for 5 min each in 0.1X SSC at 60°C. Coverslips were then counterstained with DAPI for 2 min, washed in 2X SSC and mounted in Slowfade Gold antifade (Invitrogen S36937).

**RNA FISH**

The probe for RNA FISH for HOXA locus was prepared from the BAC clone (RP11-1132K14) by nick translation. The probe was resuspended in 10µl of deionized FA and mixed with an equal volume of 2X hybridization mix (10% Dextran Sulphate and 0.1 mg salmon Sperm DNA in 2X

SSC solution, pH-7.4) containing 2mM vanadyl ribonucleoside complex (VRC), and incubated on ice for 30 min. Cells were washed thrice in ice-cold 1X PBS/ 2mM VRC (5 min each) and treated with CSK buffer/2mM VRC on ice for 5 min followed by fixation using 4 % PFA/ 2mM VRC (7 min RT). The cells were incubated with 70% ethanol at −20°C for 60 mins (or stored in 70% ethanol at −20°C until further use) followed by washes in ethanol series (70–90–100% ethanol) and air-dried. Cells were hybridized with HOXA RNA probe by incubating at 37°C overnight, followed by washes with 50% FA/2X SSC (with 2mM VRC) and 2XSSC (with 2mM VRC, pH 7.2-7.4) at 42°C (three washes each of 5 min each). The cells were mounted using Antifade followed by imaging on a confocal microscope.

**Chromatin Immunoprecipitation (ChIP)-qPCR and ChIP-sequencing**

ChIP was performed as described previously(Labade et al. 2016). In brief, Cells (~1.0 X107) were cross-linked using 1% formaldehyde for 10 min at RT. Cross-linking was quenched using 150 mM glycine, and cells were lysed in 1 ml swelling buffer (25 mM HEPES pH 8.0, 1.5 mM MgCl2, 10 mM KCl, 0.1% NP-40, 1X PIC, and nuclei were recovered by centrifugation at 2000 rpm. Fixed nuclei were re-suspended in 1 ml sonication buffer (50 mM HEPES pH 8.0, 140 mM NaCl, 1 mM EDTA, 1% Triton-X-100, 0.1% Na-deoxycholate and 0.1% SDS) supplemented with protease inhibitor cocktail and sonicated using Bioruptor Tween sonicator (Diagenode) to generate fragment sizes of ~100-500 bp. The supernatant was pre-cleared using protein A Dynabeads (Invitrogen), 1 hour at 4˚C. The amount of DNA was estimated using Nanodrop 2000. Nup93 antibody validated according to ENCODE guidelines (~2 μg) was added to ~100 μg of chromatin sample and diluted to ~1 ml in sonication buffer and incubated overnight at 4˚C [91]. CTCF antibody is a ChIP-grade antibody from Millipore (07-729, Lot-2375606). IP complexes were captured using protein-A Dynabeads (pre-blocked with 0.5 % BSA/1X PBS) and washed 3 times (at 4˚C, 11 rpm on end to end rotor) each with sonication buffer, Wash buffer-1 (50 mM HEPES pH 8.0, 500 mM NaCl, 1 mM EDTA, 1% Triton X-100, 0.1% Na-deoxycholate and 0.1% SDS), wash buffer-2 (20 mM Tris HCL pH 8.0, 1 mM EDTA, 0.5% NP-40, 250 mM LiCl, 0.5% Na-deoxycholate and 1X PIC) and TE buffer. Immunoprecipitated chromatin was eluted twice in 200 µl of elution buffer (50 mM Tris pH 8.0, 1 mM EDTA, 1% SDS, 50 mM $NaHCO_3$) at 65˚C for 10 min. Input and IP fractions were treated with 20 µg RNase A for 1h at 42˚C followed by 40 µg Proteinase K for 1 h at 65˚C. Reverse crosslinking was performed at 65˚C overnight. DNA was extracted with phenol: chloroform: isoamyl alcohol (25:24:1), and ethanol precipitated using 3M sodium acetate (pH 5.2) and 2 µg of glycogen. DNA samples were washed with 70% ethanol and re-suspended in 10 µl of nuclease-free water.

**Microscopy and Image analysis**

Image acquisition was performed on a Zeiss LSM710 confocal microscope with a 63X Plan-Apo1.4 NA oil immersion objective with 405 nm, 488 nm, and 561 nm laser at 1 to 2.5 digital zoom. Acquisition of Z-stacked images (voxel size of 0.105µm × 0.105µm × 0.34µm) was at 512 × 512 pixels per frame using 8-bit pixel depth for each channel and a pinhole size of 0.7µm (1 arbitrary unit [AU]). The line averaging was set to 4.0, and images were collected sequentially in a three-channel mode.

Distances of gene loci from the nuclear periphery were measured (in µm) in 3D using the boundary of the DAPI signal as a marker of the nuclear periphery (Shachar et al. 2015). For quantification, confocal images were loaded into the object analysis tool of the Huygens Professional software (Version 18.10.0p2 64b, built Nov 20, 2018). This tool performs surface rendering according to the threshold segmentation of different group of voxels that are separated from the background into a 3D object. 3D reconstruction was performed using surface rendering for the nucleus (blue channel), HOXA gene locus (red channel), CT7 (green channel). Comparable threshold and seeding levels were used for surface rendering for all images. The nucleus was selected as an anchor, and the shortest distance of each gene locus (Object) from the nuclear periphery was measured using the ‘analyze object tool’ in Huygens professional software (Version 18.10.0p2 64b, built Nov 20, 2018).

**ChIP-sequencing**

Chromatin immunoprecipitation of Nup93 associated DNA was performed as mentioned in the materials and methods section. Nup93 ChIP-DNA (Two biological replicates) and Input DNA was outsourced for high throughput sequencing to Genotypic Technology, Bangalore, India. ChIP-Seq libraries for sequencing were constructed according to the NEXTflex™ ChIP-Seq library protocol outlined in NEXTflex™ ChIP-Seq Kit - 5143-01. Briefly, DNA was subjected to a series of enzymatic reactions that repair frayed ends, phosphorylate the fragments, add a single nucleotide A overhang and ligate adaptors (NEXTflex ChIP Barcodes-48 kit). The libraries were enriched using PCR (5 cycles), fragments were size selected using a 2 % Low melting agarose gel and purified using MinElute Gel Extraction Kit (QIAGEN). The libraries were further enriched using PCR (14 cycles), post PCR cleanup was performed using Agencourt AMPURE XP beads (Beckman Coulter #A63881). The prepared libraries were quantified using Qubit fluorometer and validated for quality by running an aliquot on High Sensitivity Bioanalyzer Chip (Agilent).

**ChIP-seq analysis**

Quality control for all Illumina sequence reads was performed using a FastQC tool (Version 0.72) (Afgan et al. 2018)**.** All quality-controlled reads were mapped against the reference Human genome (hg19) using Bowtie. Reads that aligned uniquely to a single genomic location were used for further analysis. Correlation between two biological replicates of ChIP-seq was determined using ‘multiBamSummary’ and ‘plotCorelation’ tool on GALAXY server. Aligned BAM files were visualized using genome browser within EaSeq platform as well as on UCSC genome browser. Since both replicates were significantly correlated with one another only onc replicate was processed for further analysis.

**Peak calling**

MACS peak (version 1.4.2 20120305) calling algorithm was used to find peaks representing likely binding sites for Nup93. Parameters used for peak calling are as follows

### effective genome size = $2.70e^{+09}$

### band width = 300

### model fold = 10,20

### p-value cutoff = $1.00e^{-05}$

### Large dataset will be scaled towards smaller dataset.

### Range for calculating regional lambda is: 1000 bps and 10000 bps

Significant peaks identified by MACS were further filtered and ranked based on p-value. Wiggle files generated using MACS were used for visualization on the genome browser. Statistically significant peaks obtained from MACS peak caller were used for peak annotation. Peaks were annotated using CEAS (cis-regulatory element annotation system) (Shin et al. 2009). CEAS annotate each peak summit with the genomic annotations in the following four categories: (a) promoters, (b) bidirectional promoters, (c) downstream of a gene, and (d) gene bodies (3'UTRs, 5'UTRs, coding exons, and introns). Promoter regions were defined as -1 kb, -2kb and -3kb upstream of the TSS. Downstream regions were defined as +1kb, +2kb and +3kb downstream of the Transcription End Site (TES). Each peak was annotated with the closest gene found within a 500 kb window from the center of the summit. We also performed independent gene annotation using EaSeq platform (Lerdrup et al. 2016).

**Gene ontology and Motif enrichment analysis:**

Gene ontology analysis of Nup93 associated genes was performed using DAVID (https://david.ncifcrf.gov/home.jsp). We also performed KEGG pathway analysis of Nup93 associated genes (Kyoto Encyclopedia of Genes and Genomes, http://www.genome.jp/kegg/). For Motif analysis, Nup93 associated FASTA sequences were submitted as an input to MEME ChIP and motifs analysis was performed with default parameters against HOCOMOCO Human (v11 FULL) database (Machanick and Bailey 2011; Bailey et al. 2009). Out of 20 motifs identified by MEME-ChIP, top 6 motifs were selected for further analysis. We performed TOMTOM (Tomtom compares one or more motifs against a database of known motifs) analysis on to 6 motifs to identify similar motifs from HOCOMOCO Human (v11 FULL) database (Bailey et al. 2009). To count the number of high probability CTCF motifs within Nup93 binding sequences we used MEME motif finding software [FIMO version 5.0.5, (Release date: Mon Mar 18 20:12:19 2019 -0700)]

**Transcription factors and Histone mark enrichment analysis**

We performed transcription factor enrichment analysis of Nup93 associated peaks using ReMap (Regulatory Map of TF Binding Sites)(Chèneby et al. 2018). 404 Nup93 binding peaks (.bed) were submitted to ReMap analysis (with 10 % Minimum overlap) against the ReMap catalog of transcription factor binding peaks. ChIP-atlas database (http://chip-atlas.org/) was used to look at the enrichment of transcription factors and histone marks specifically in DLD-1 cells. We downloaded 'BAM' and 'bigwig' files for CTCF (DRX013180), H3K4me1(DRX013183), H3K4me3 (DRX013175) H3K27ac (DRX013177), H3K27me3 (DRX013182) and H3K36me3 (DRX013178) from DLD-1 cells. Correlation plots and heatmaps were generated using EaSeq platform (Lerdrup et al. 2016).

**Table S1: List of siRNA sequences, RT-qPCR primers and ChIP-qPCR primers**

| | |
|---|---|
| **siRNA sequences** | |
| Nup93 | 5′-AGAGTGAAGTGGCGGACAA-3′ |
| Nup188 | 5′-GCCTTTCTGCGCTTGATCACCACCC-3′ |
| Nup205 | 5′-AGAUGGUGAAGGAGGAAUAUU-3′ |
| CTCF | 5′-CAAGAAGCGGAGAGGACGA-3 |
| Nup98 | 5'-TGTCAGACCCTAAGAAGAA-3 |
| LacZ | 5'-CGUACGCGGAAUACUUCGA-3' |
| **List of RT-qPCR primers** | |
| Nup93 | F-AGAAGACGCCCTTGACTTTAC<br>R-GATATAAATTTGCCGCGCATAGG |
| Nup188 | F-CTGGGCAATCAGCAGGATATAA<br>R-AATGATCCCAAGGCCAGAAG |
| Nup205 | F-GACCCTAGAACTCAGTCCAGA<br>R-CTGTGACACCAGCGTAAGAA |
| CTCF | F-CGTTACTGTGATGCTGTGTTTC<br>R-TCATGTGCCTCTCCTGTCTA |
| HOXA1 | F-CGTTAAATCAGGAAGCAGACCC<br>R-GTAGCCGTACTCTCCAACTTTC |
| HOXA5 | F-CTGCACATAAGTCATGACAACATAG<br>R-GGTCAGGTAACGGTTGAAGT |
| HOXA9 | F-AAAAGCGGTGCCCCTATACA<br>R-CGGTCCCTGGTGAGGTACAT |
| HOXA13 | F-AGCGCGTGCCTTATACCAAG<br>R-GCCGCTCAGAGAGATTCGT |
| Nup98 | F-CCATCTATGGATGACCTTGCTAA<br>R-TCCGACCAATAGTGAAATCAGAG |
| Oct4 | F-AGCAAAACCCGGAGGAGT<br>R-CCACATCGGCCTGTGTATATC |
| Sox2 | F-AGACGCTCATGAAGAAGGATAAGT<br>R-CTGCGAGTAGGACATGCTGTAG |
| Nanog | F-GAAATCTAAGAGGTGGCAGAAAAA<br>R-GCAGAGATTCCTCTCCACAGTTAT |
| **List of ChIP-qPCR primers** | |
| HOXA1 promoter | F-CGCTCTTCCCCCTCCATT<br>R-ACCGTTCAATGAAAGATGAACTG |
| HOXA5 Promoter | F-TGTATGGAATTTGACCTCGC<br>R-CAACAACTTTATTTCCCCCG |
| CBS1 | F-GCGGTCGTTTGTGCGTCTAT<br>R-CCTCCACCCAACTCCCCTATT |
| CBS2 | F-AATCCCAAAGCCAGAGTGTT<br>R-GCTGGACGCCGTTATAGACT |
| CBS4 | F-GGGTGAGTCCCCTTTTTCTGTT<br>R-GCGCACTTCCGATCAATGTC |
| CBS5 | F-GGAGGATGCTCGCAGGACAC<br>R-GGGCGGGAAGGGGAGTAT |
| HOXA1 upstream region (~2700bp upstream TSS) | F-CTGAAAGAGGCGTTTTGAGC<br>R-GGAGCTGGTCTCTTTCAACG |
| HOXA1 downstream region (~1750bp downstream of TES) | F-ATGAATGCAGTGATGGGTCA<br>R-AACCAAGAGGGGAGAGGAAA |

#### SUPPLEMENTARY FIGURE LEGENDS

**Supplementary Figure 1.** Genome-wide occupancy of Nup93. **(A)** Scatter plot showing Pearson's correlation coefficient (r) between signal intensities ($Log_2$ read counts in 10 kb bins) of replicate1 and replicate2 of Nup93 ChIP-seq data sets from DLD-1 cells. Two replicates of Nup93 ChIP-seq shows a strong correlation with each other (Pearson's correlation coefficient=0.92). **(B)** UCSC genome browser view of LMNA gene locus showing overlapping Nup93 ChIP-seq peaks from two biological replicates. **(C)** Histogram showing enrichment of Nup93 peaks around (± 3Kb) transcription start sites (TSS) of annotated genes. **(D)** Ideogram representing cytogenetic positions of Nup93 peaks on all chromosomes. **(E)** A plot representing % Nup93 occupancy as against gene density for all human chromosomes. **(F)** A plot representing % Nup93 occupancy as against chromosome size for all human chromosomes. Nup93 occupancy does not correlate with gene density or chromosomal size. % occupancy = [(number of peaks per chromosome/ total number of peaks)]

**Supplementary Figure 2.** Genome-wide occupancy of Nup93. **(A)** Enrichment analysis of Nup93 binding sites. KEGG pathway categories of Nup93 associated genes. P-value represents the significance of enrichment. **(B)** Top DNA-binding motifs enriched among binding sites of Nup93 as revealed by MEME-ChIP analysis, with corresponding p values. **(C)** Transcription factor enrichment analysis of Nup93 binding peaks using ReMap (Chèneby et al. 2018). Table representing the top three transcription factors enriched on Nup93 binding sites (sorted according to observed overlap). **(D)** Signal tracks representing an enrichment of Nup93 and CTCF on the promoter (dotted black box) of the HOXA1 gene in DLD-1 cells. The area under the peak represents the number of reads mapped in that genomic region. The *Y-axis* represents normalized peak intensity. The blue bar underneath the signal tracks indicates the HOXA1 gene.

**Supplementary Figure 3.** Characterization of NT2/D1 differentiation. **(A)** A graph showing fold change ($2^{-\Delta\Delta Ct}$) in transcript levels of Pax6 normalized to Day 0. Data from N=3 independent biological replicates for Day 0,2, 4, and 8. N=2 for Day 21. *Y-axis*: Fold change ($2^{-\Delta\Delta Ct}$) in expression levels normalized to Day 0. Error bars: SEM. **(B)** Representative images of cell cycle profile of NT2/D1 cells during RA treatment across days. **(C)** Cell cycle profile analysis of NT2/D1 cells upon RA treatment on Day 0, 2, 4 & 8. The graph represents % distribution profiles of cells in different cell cycle stages. Data from N=2 independent biological replicates. Error bars: SD. **(D)** Densitometry quantification of western blots from Fig. 2E, showing Nup93, Nup188, Nup205 and CTCF levels on Day 0, 2, 4, & 8. Protein levels normalized to respective GAPDH. Data from N=3 independent biological replicates. Error bars: SEM. Student's t-test. (* = p<0.05, **= p<0.01). **(E)** Quantification of HOXA RNA-FISH foci on Day 0 (n=80), Day 2 (n=72), Day 4 (n=84), and Day 8 (n=149) upon RA treatment, n: number of nuclei. Y-axis represents % nuclei showing 0-4 copies of active HOXA RNA foci during differentiation indicative of actively transcribing HOXA genes upon RA treatment. Data from N=2 independent biological replicates, error bar=SD

**Supplementary Figure 4.** Characterization of HOXA FISH probe. **(A)** Schematic representation of HOXA gene locus showing positions of BAC clones used for HOXA FISH probe preparation. **(B)** Representative image showing metaphase spread of NT2/D1 cells. Scale bar= 10μm. **(C)** Histogram showing quantification of chromosome number in NT2/D1 cells. *Y-axis*: % of cells with chromosome counts. **(D)** Representative 2D FISH image showing DAPI (blue), Chromosome 7 (Green) and HOXA locus (red) in NT2/D1 metaphase spreads. NT2/D1 cells harbor 2 full copies of Chr. 7 and 2 truncated 'p' arms. 4 copies of HOXA gene locus can be detected. Representative data from a single experiment from 15 different metaphase spreads. **(E)** Enlarged images of four copies of Chr.7 (Two full and two truncated copies) with four HOXA gene loci (red).

**Supplementary Figure 5.** Nup93 and CTCF regulate HOXA gene expression in an antagonistic manner in undifferentiated embryonal carcinoma cells. **(A)** Densitometry quantification of western blots from Fig. 4C showing levels Nup93, CTCF and Oct4 in UT, siLacZ, Nup93 Kd, CTCF KD, Nup93 and CTCF Kd and Nup98 Kd. Protein levels normalized to respective GAPDH. Data from N=2 independent biological replicates. Error bars: SD. **(B)** Cell cycle profile analysis of NT2/D1 cells upon siLacZ, Nup93 Kd, Nup188 Kd, Nup205 Kd, CTCF Kd, and Nup98 Kd. The graph represents % distribution profiles of cells in different cell cycle stages. Data from N=1 single experiment. **(C)** Representative images (maximum intensity projections) of Poly-A RNA FISH in siLacZ (n=30), Nup93 Kd (n=30) and Nup98 Kd (n=28) cells. DAPI (blue) and Oligo dT probe (green). Scale bar = 10µm. White *arrowhead* indicates nuclear bloc of PolyA in Nup98 Kd cells. **(D)** Scatter plot showing Nuclear/ cytoplasmic ratio of Poly-A signals in siLacZ (median = 0.72), Nup93 Kd (median = 0.84) and Nup98 Kd (median = 0.89). Data from a single experiment. The *horizontal bar* represents the median with interquartile range. Significance calculated using Mann-Whitney U test between siLacZ and knockdown samples (***= p<0.001). **(E, F)** Biological replicate2 for RT-qPCR data shown in (Figure 4E, F). A graph representing fold change ($2^{-\Delta\Delta Ct}$) in transcript levels of HOXA1, HOXA 5 and HOXA9 upon **(E)** Nup93 Kd and **(F)** CTCF knockdown followed by RA treatment. Data from N=1 experiment which includes n=2 independent technical replicates. Error bars: SEM. **(G, H)** Effect of Nup93 and CTCF Kd on HOXA1 expression at early time points upon RA treatment. A graph representing fold change ($2^{-\Delta\Delta Ct}$) in transcript levels (X-axis) of HOXA1 from 30min to 4h of RA treatment (X-axis).

**Supplementary Figure 6.** Biological replicate 2 for ChIP-qPCR data shown in Figure 6. **(A-D)** CTCF ChIP-qPCR shows a gradual decrease in CTCF occupancy on its conserved binding sites **(A)** CBS1, **(B)** CBS2, **(C)** CBS4 and **(D)** CBS5 upon RA treatment during differentiation. **(E, F)** Nup93 ChIP-qPCR shows a gradual decrease in its occupancy on **(E)** HOXA1 and **(F)**

HOXA5 promoters from Day 0 to Day 8 and in response to RA treatment. *Y-axis*: immunoprecipitated DNA relative to 1% input, N=1 data from one experiment that include a total of three technical replicates, error bar=SEM.

**Supplementary Figure 7.** ChIP-qPCR controls experiments for data represented in Fig. 6 (Biological replicate1). **(A, B)** ChIP-qPCR showing Nup93 and CTCF enrichment on a control region ~3Kb **(A)** upstream and **(B)** downstream of HOXA1 promoter during differentiation. **(C-J)** ChIP-qPCR showing enrichment of antibody control (PanH3) on **(C)** HOXA1 promoter, **(D)** HOXA5 promoter, **(E)** CBS1, **(F)** CBS2, **(G)** CBS4, **(H)** CBS5, **(I)** HOXA1 upstream region and **(J)** HOXA1 downstream region during RA treatment. *Y-axis*: Immunoprecipitated DNA with respect to 1% input.

**Supplementary Figure 8.** ChIP-qPCR controls experiments for data represented in Fig. 6 (Biological replicate-2). **(A, B)** ChIP-qPCR control experiments for data represented in Figure S7. ChIP-qPCR showing Nup93 and CTCF enrichment on a control region ~3Kb **(A)** upstream and **(B)** downstream of HOXA1 promoter during differentiation. *(C-J)* ChIP-qPCR showing enrichment of antibody control (PanH3) on **(C)** HOXA1 promoter, **(D)** HOXA5 promoter, **(E)** CBS1, **(F)** CBS2, **(G)** CBS4, **(H)** CBS5, **(I)** HOXA1 upstream control and **(J)** HOXA1 downstream control during RA treatment. *Y-axis*: Immunoprecipitated DNA with respect to 1% input.

#### Supplementary Figure S1

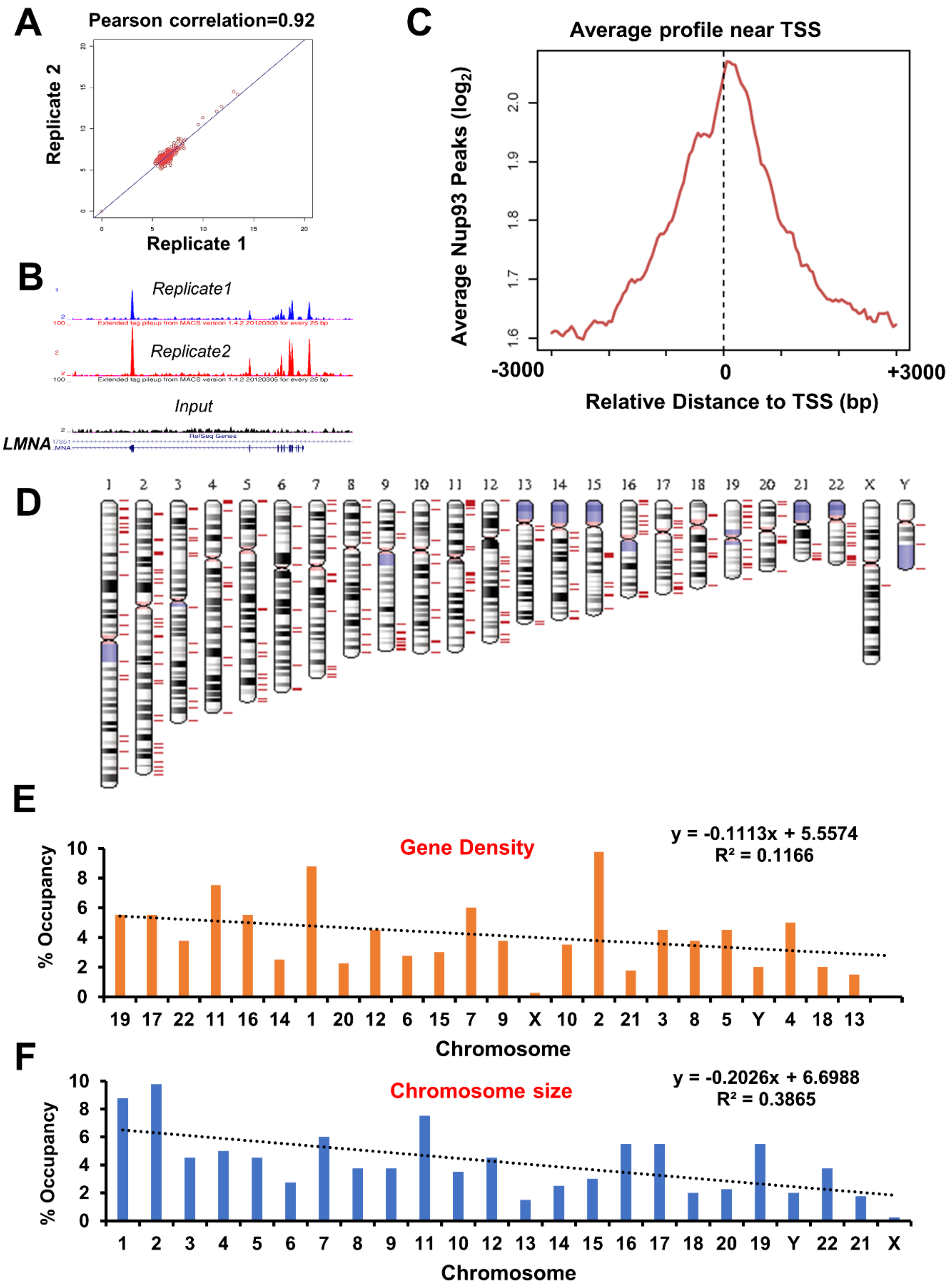

#### Supplementary Figure S2

**A**

| KEGG Pathways | -log ( p-value) |
|---|---|
| Hippo signaling pathway | 20.72 |
| Signaling pathways regulating pluripotency of stem cells | 12.15 |
| Basal cell carcinoma | 11.26 |
| Wnt signaling pathway | 10.02 |
| Colorectal cancer | 7.72 |
| Oxytocin signaling pathway | 7.13 |

**B**

| CONSENSUS MOTIF | E-VALUE | SIMILAR_MOTIF |
|---|---|---|
| 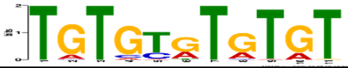 | 1.20E-54 | EGR3 |
| 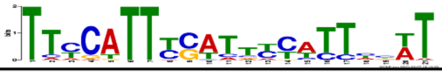 | 5.60E-36 | ZN394 |
| 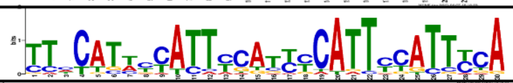 | 2.80E-28 | ZN502 |
| 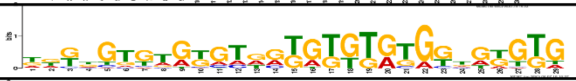 | 3.50E-24 | RREB1 |
| 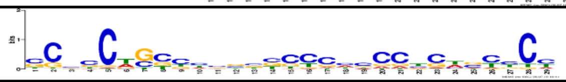 | 6.30E-22 | MAZ |
| 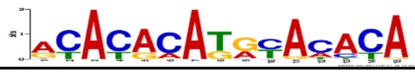 | 3.70E-17 | FOXG1 |

**C**

| Transcription Factor | Observed Overlap | $-log_{10}$(E-value) |
|---|---|---|
| CTCF | 324/404 | 84.244 |
| MYC | 300/404 | 73.447 |
| AR | 278/404 | 46.275 |

**D**

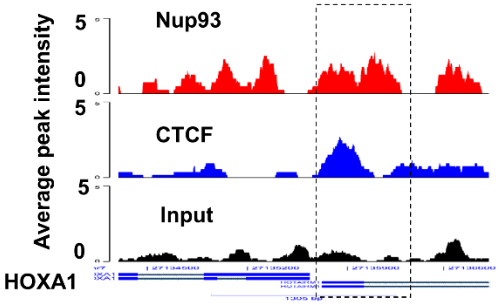

#### Supplementary Figure S3

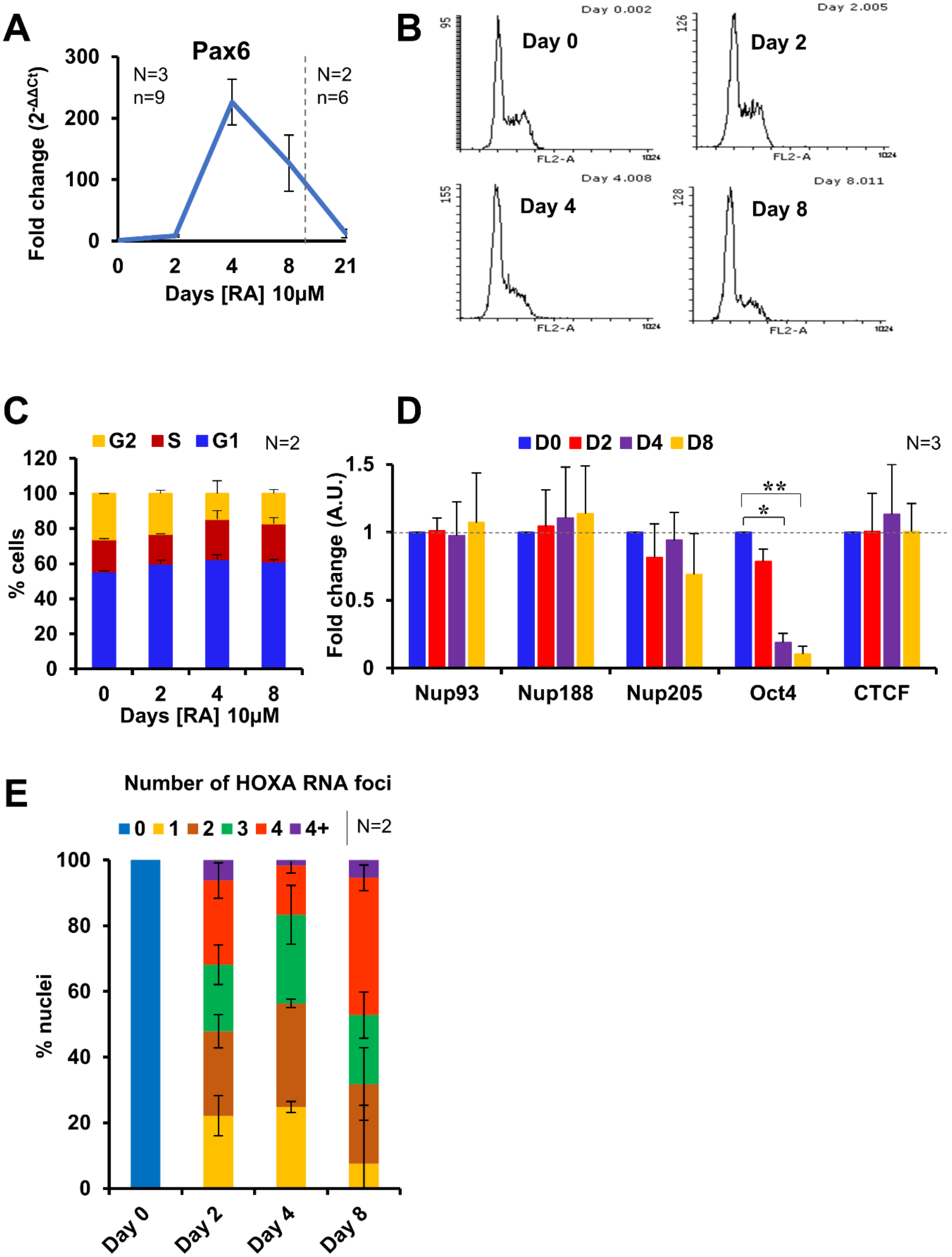

Supplementary Figure S4

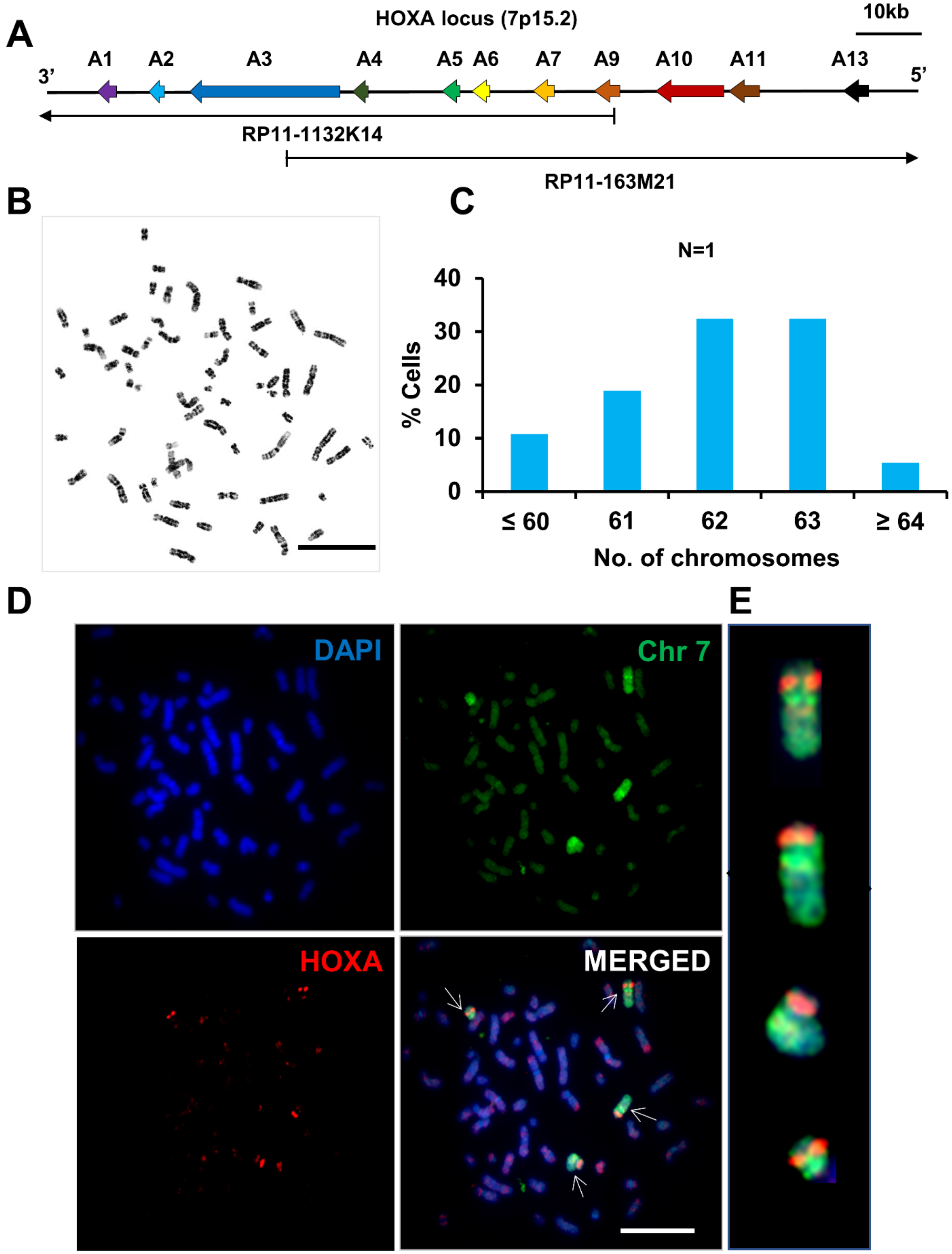

#### Supplementary Figure S5

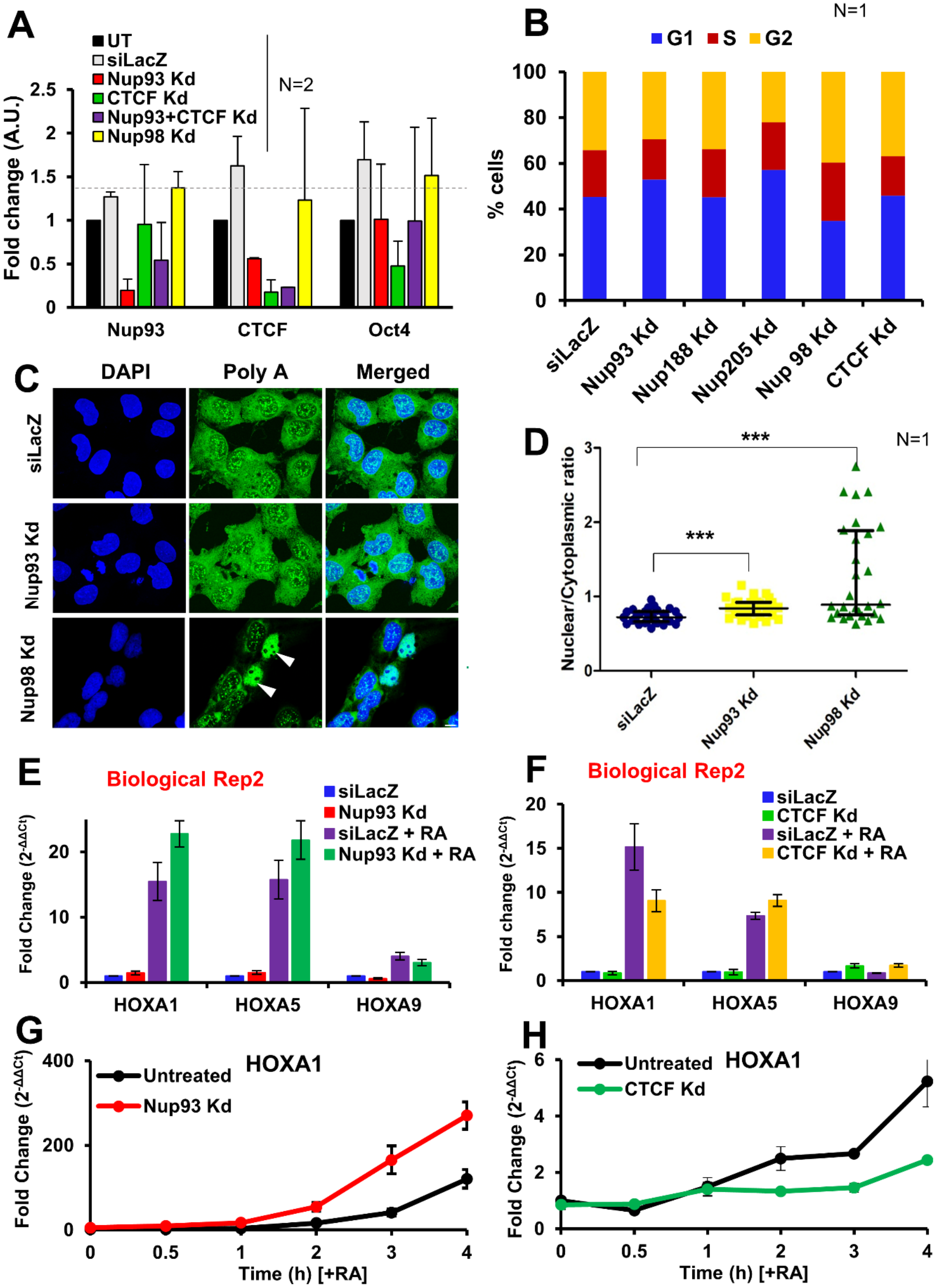

Supplementary Figure S6

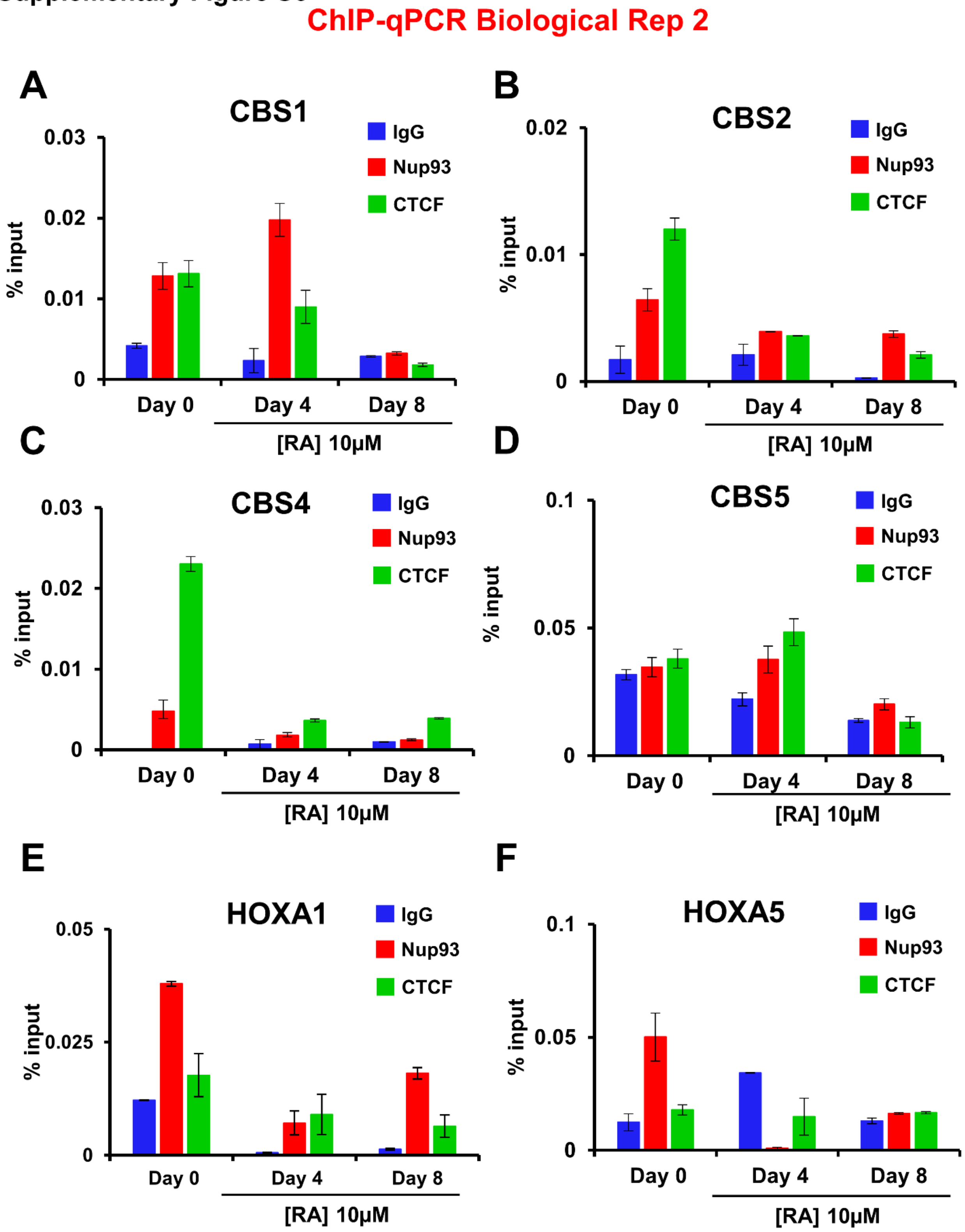

**Supplementary Figure S7**

#### ChIP-qPCR controls- Biological Rep1

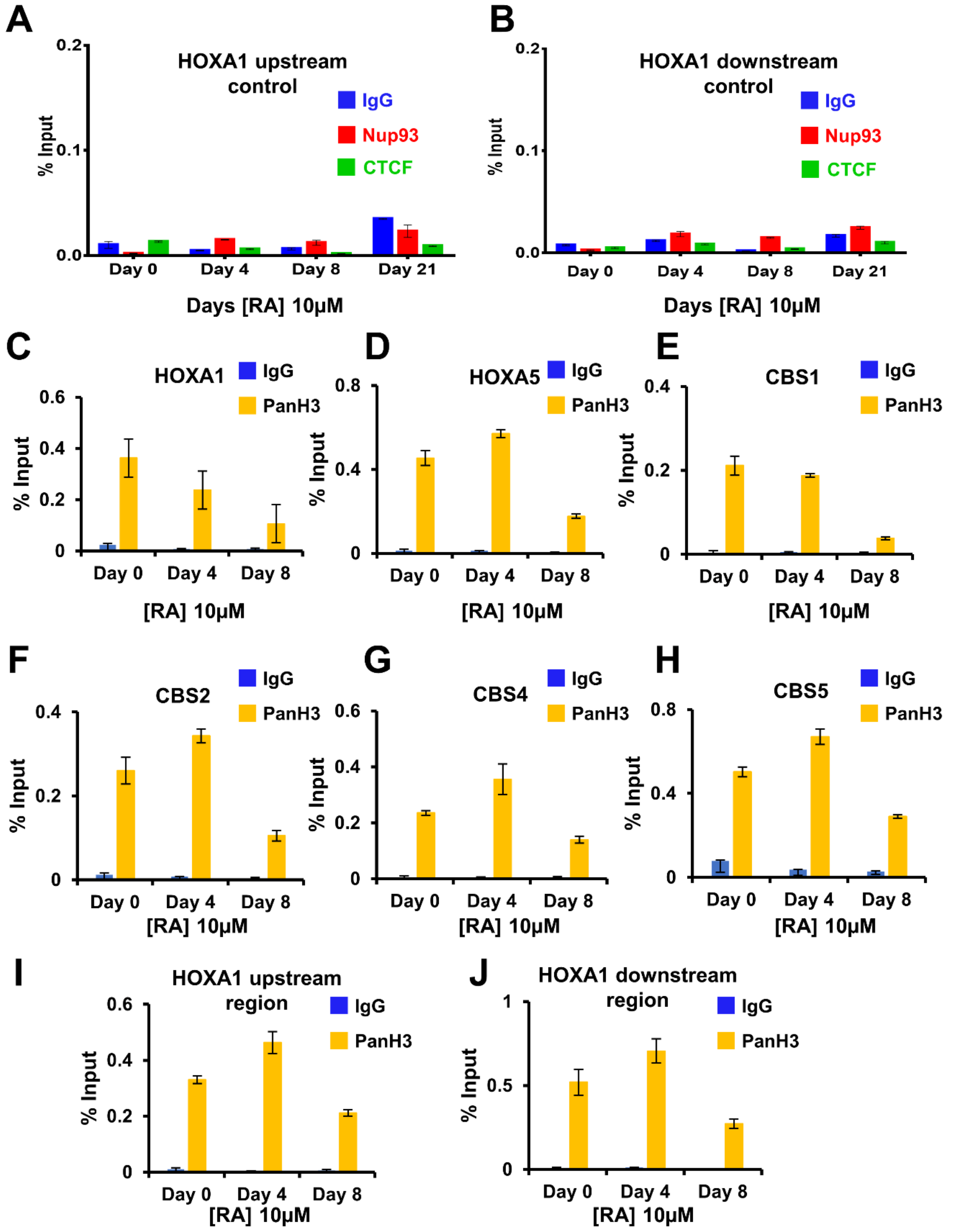

Supplementary Figure S8

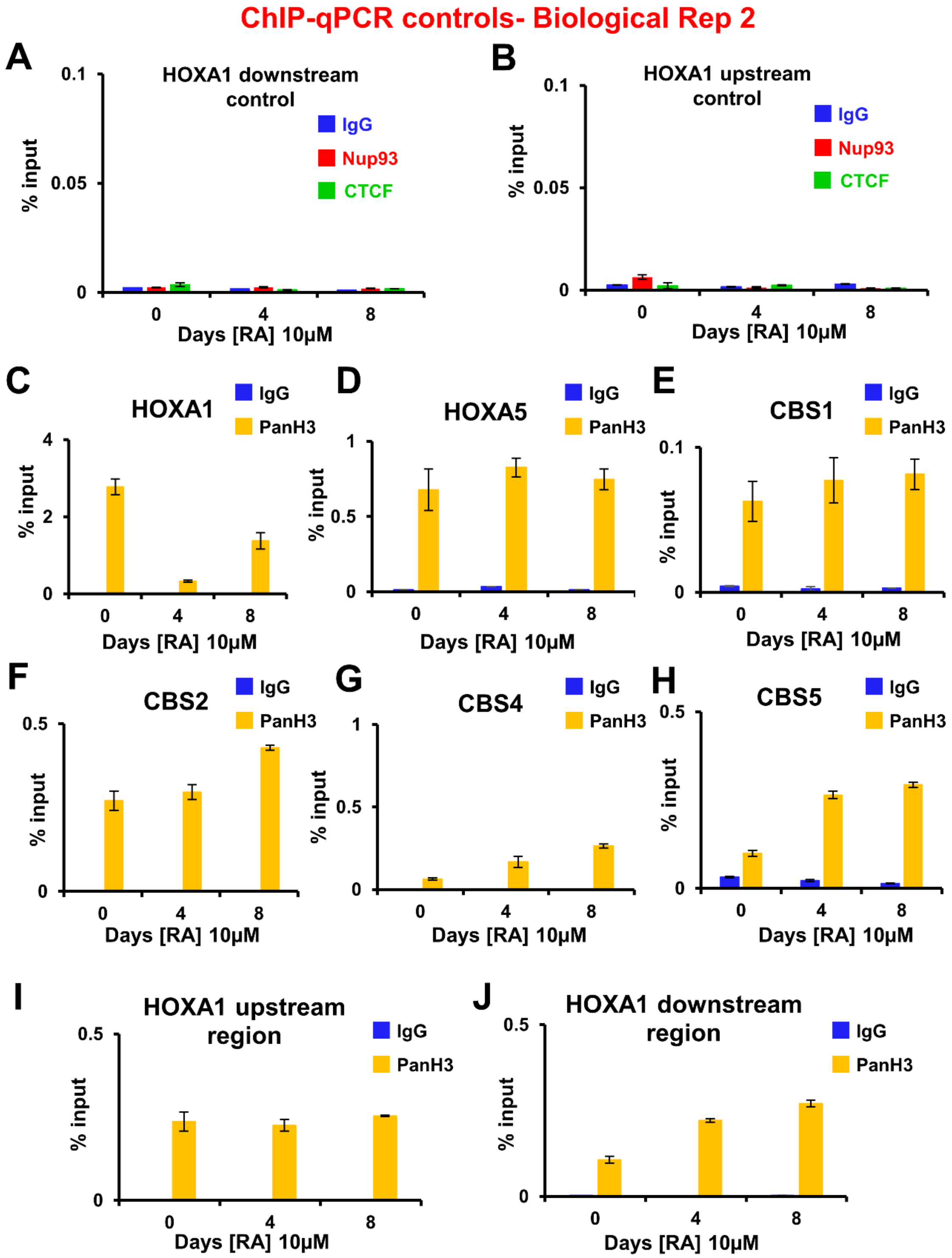
